# Parp7 generates an ADP-ribosyl degron that controls negative feedback of androgen signaling

**DOI:** 10.1101/2024.12.21.629908

**Authors:** Krzysztof Wierbiłowicz, Chun-Song Yang, Ahmed Almaghasilah, Patryk A. Wesołowski, Philipp Pracht, Natalia M. Dworak, Jack Masur, Sven Wijngaarden, Dmitri V. Filippov, David J. Wales, Joshua B. Kelley, Aakrosh Ratan, Bryce M. Paschal

**Affiliations:** Department of Biochemistry and Molecular Genetics, University of Virginia School of Medicine, PO Box 800733, Charlottesville, VA 22908, USA; Center for Public Health Genomics, University of Virginia School of Medicine, Charlottesville, Virginia; Department of Molecular and Biomedical Sciences, University of Maine, Orono, ME, USA; Yusuf Hamied Department of Chemistry, University of Cambridge, Lensfield Road, Cambridge, CB2 1EW, U.K; Advanced Microscopy Facility, University of Virginia School of Medicine, Charlottesville, VA 22908, USA; University of Virginia Comprehensive Cancer Center, Charlottesville, VA 22903, USA; Leiden Institute of Chemistry, Leiden University, Einsteinweg 55, Leiden 2333 CC, The Netherlands

**Keywords:** E3 ligase, androgen, ubiquitin, proteasome, DTX2, PARP, ADP-ribosylation, RBN2397, PARP7, negative feedback

## Abstract

The androgen receptor (AR) tranduces the effects of circulating and tumor-derived androgens to the nucleus through ligand-induced changes in protein conformation, localization, and engagement with chromatin binding sites. Understanding these events and their integration with signal transduction is critical for defining how AR drives prostate cancer and unveiling pathway features that are amenable to therapeutic intervention. Here, we describe a novel post-transcriptional mechanism that controls AR protein levels on chromatin and associated gene output which is based on a highly selective, inducible degradation mechanism. We find that the mono-ADP-ribosyltransferase PARP7 generates an ADP-ribosyl degron on a single cysteine within the DNA binding domain of AR, which is then recognized by the ADP-ribose reader domain in the ubiquitin E3 ligase DTX2 and degraded by the proteasome. Mathematical modeling of the pathway suggested that PARP7 ADP-ribosylates chromatin-bound AR, a prediction that was validated in cells using an AR mutant that undergoes nuclear import but fails to bind DNA. Lysine-independent, non-conventional ubiquitin conjugation to ADP-ribosyl-cysteine and AR degradation by the proteasome forms the basis of a negative feedback loop that regulates specific modules of AR target genes. Our data expand the repertoire of mono-ADP-ribosyltransferase enzymes to include gene regulation based on highly selective protein degradation.

**One Sentence Summary:** PARP7 mono-ADP-ribosylates the androgen receptor on Cys620 to mark the androgen receptor for ubiquitin conjugation by an E3 ligase with ADP-ribose reader function, resulting in in negative feedback of AR-dependent gene expression.

## INTRODUCTION

The poly-ADP-ribosyltransferase (PARP) family of enzymes use NAD^+^ as a cofactor to conjugate ADP-ribose to substrates. Most PARP family members mediate mono-ADP-ribosylation, though a subset including PARP1 and PARP2 generate poly-ADP-ribose chains (PAR)(*1*). As the founding member of the PARP family, PARP1 has been studied extensively, especially in the context of DNA repair and cancer.

PARP1/2 inhibitors are FDA-approved for the treatment of several cancers, with the greatest activity against tumors with homologous recombination deficiency(*2*). For the PARP family members that mediate mono-ADP-ribosylation, major knowledge gaps limit our understanding of associated biology and therapeutic opportunities. What is clear is that the mono-ADP-ribosyltransferases participate in a variety of biological pathways that impact major cellular events, including gene expression, cell identity, and immune function, among others(*3, 4*).

The exact mechanisms by which ADP-ribosylation regulates protein function are also under active investigation. PARP1 auto-modification promotes its release from chromatin, and PARP1-generated poly-ADP-ribose chains provide scaffolds at damage sites that concentrate DNA repair factors(*5, 6*). This observation indicates that ADP-ribosylation, like several other post-translational modifications, including those occurring in the nucleus, can regulate protein dynamics. Protein modules with three-dimensional structures known to read ADP-ribosylation are the macrodomain, Deltex C-terminal (DTC) domain, Tryptophan-Tryptophan-Glutamate (WWE) domain, and the PAR-binding zinc-finger (PBZ) domain(*7, 8*). As these structural modules occur in a variety of proteins, including some PARP enzymes, ADP-ribose readers can act as ADP-ribosylation effectors that promote PARP- and substrate-specific outcomes.

PARP activity in the nucleus has also been shown to intersect with processes associated with transcription. Multiple studies have linked PARP1 and PARP2 to transcription through effects on chromatin structure(*9*). Another example of PARP regulation of transcriptional output involves the mono-ADP-ribosyltransferase 2,3,7,8-Tetrachlorodibenzodioxin-inducible PARP (TIPARP), also known as PARP7. Induction of PARP7 by the aryl hydrocarbon receptor (AHR) is part of a mechanism that limits the AHR-mediated xenobiotic response, which otherwise promotes hepatotoxicity(*10, 11*). Our group identified PARP7 in the context of androgen signaling through AR(*12*). AR directly induces PARP7 expression, which mono-ADP-ribosylates AR on Cys residues; these ADP-ribosyl-Cys sites are read by PARP9 macrodomains in the DTX3L/PARP9 complex and results in the assembly of a DTX3L/PARP9-AR complex(*12*). Thus, PARP7 induction, substrate (AR) mono-ADP-ribosylation, and reading by a complex that contains an E3 ligase for histone mono-ubiquitylation(*13*) contribute to AR regulation of gene expression. A striking feature of the mechanism is that agonist-conformation-dependent AR ADP-ribosylation by PARP7 occurs on eleven cysteines(*12, 14*). While the stoichiometry of ADP-ribosylation remains to be determined, the total number of ADP-ribosyl-cysteines in AR generated by PARP7 exceeds the number of macrodomains in the oligomeric DTX3L/PARP9 complex(*15*). This raised the question of whether ADP-ribosylation sites in AR might engage with other reader proteins that affect transcription.

Here, we use gene expression analysis to show that PARP7 activity has a profound effect on androgen signaling by regulating the expression of distinct modules of AR target genes through negative feedback. The mechanism is based on the androgen-induced degradation of AR, which is mediated by the ubiquitin (Ub) E3 ligase, DTX2. Our biochemical analysis shows that DTX2 uses an ADP-ribose reader domain to recognize and conjugate Ub to ADP-ribosyl cysteine in AR. These data, together with recent work(*16, 17*) indicates that PARP activity, reader function, and non-canonical ubiquitylation can be combined for highly selective protein degradation. In the case of androgen signaling, ADP-ribosylation and turnover of chromatin-associated AR illustrate how PARP-mediated degradation of a transcription factor can shape the transcriptome of prostate cancer cells.

## RESULTS

PARP7 ADP-ribosylation of AR results in selective recruitment of the DTX3L/PARP9 complex(*12*) PARP9 provides ADP-ribose reader function while DTX3L contributes E3 mono-ubiquitylation activity, potentially for substrates such as core histones(*18, 19*). Because depletion of DTX3L/PARP9 affects the expression of only 6% of the AR transcriptome(*12*), we hypothesized that multi-site AR ADP-ribosylation by PARP7 might contribute to AR regulation through additional mechanisms. To explore this hypothesis, we used a potent PARP7 inhibitor, RBN2397, that eliminates AR ADP-ribosylation in prostate cancer cells(*20, 21*). We queried the effects of PARP7 inhibition on androgen signaling in the vertebral metastasis-derived prostate cancer line VCaP using RNA-seq. VCaP cells were treated +R1881 (synthetic androgen) and +RBN2397 for 18 hours (hr), harvested, and processed. As expected, R1881 treatment alone affected the expression of a very large number of genes (9,453; using adjusted p-value (p-adj) < 0.001), with similar numbers of genes showing increased and decreased expression (Fig. 1A, upper plot). A comparison of R1881+RBN2397 vs. R1881 alone showed a substantial effect of PARP7 inhibition, with 7,861 genes differentially expressed using the same significance threshold (Fig. 1A, lower plot). In contrast, RBN2397 treatment alone affected only 19 genes (Supplementary Fig. 1A). When tested for overlap of differentially expressed genes (DEGs), 67% of the androgen-regulated genes were affected by PARP7 inhibition (Fig. 1B). The overlap included 688 genes that are among the top 1,000 direct AR target genes identified by CistromeGO in VCaP cells(*22*). These data show that PARP7 plays a major role in androgen signaling.

**Fig. 1:**
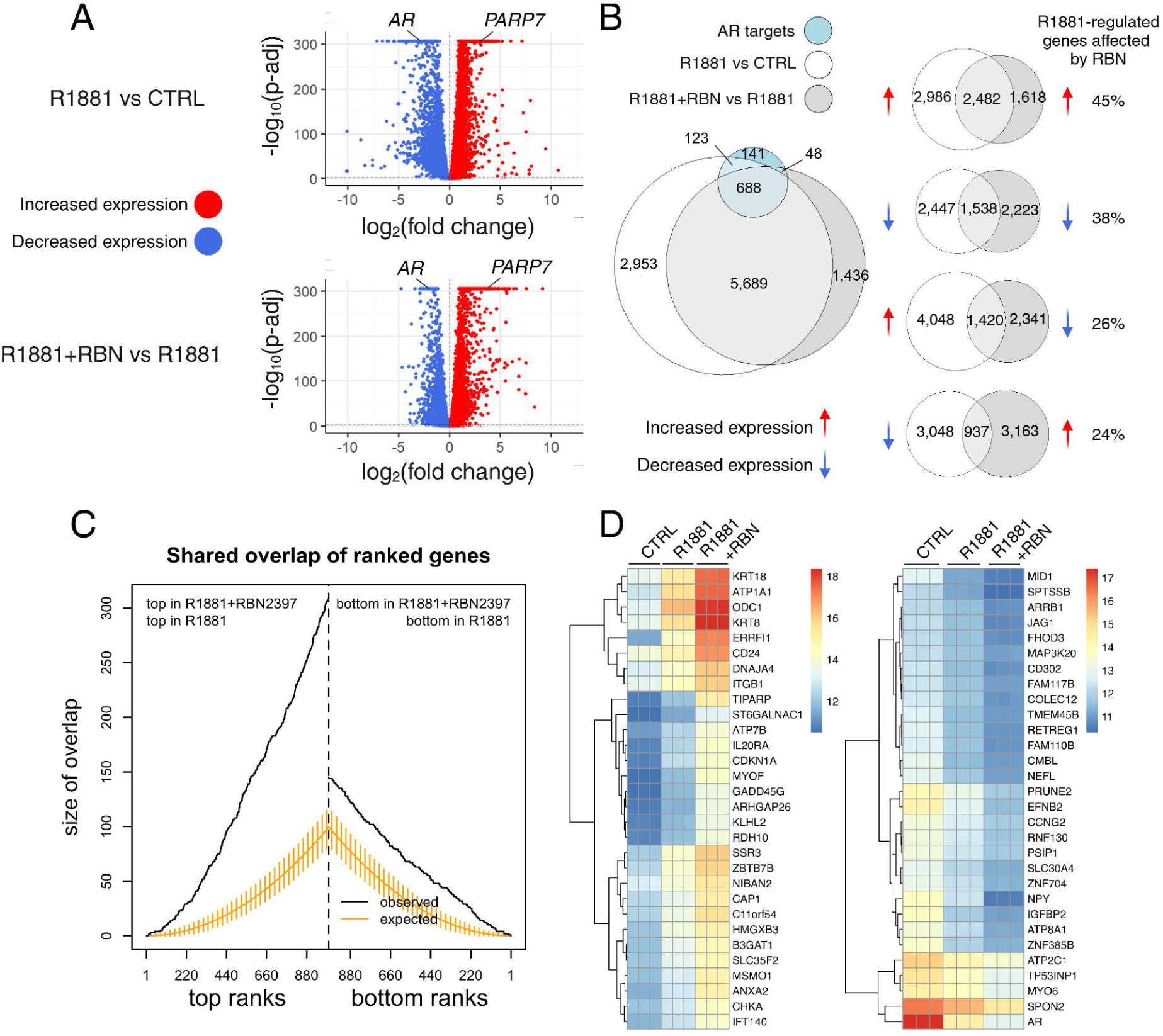
Androgen-regulated gene expression is sensitive to PARP7 inhibition by RBN2397. A, Volcano plots depicting the differential gene expression analysis in VCaP cell line among three experimental groups: untreated (CTRL), R1881-treated, and R1881+RBN2397-treated. The upper plot compares gene expression between R1881-treated and untreated samples (R1881 vs. CTRL), while the lower plot compares gene expression between R1881+RBN2397-treated and R1881-treated samples (R1881 used as a control for the comparison, R1881+RBN vs. R1881). Each dot represents a gene, with the x-axis showing the log2 fold change (Log2FC) and the y-axis representing the negative log10 of the adjusted p-value (p-adj). Genes that have significantly increased or decreased expression are highlighted in red and blue, respectively, based on the preset p-adj (0.001) threshold. AR and PARP7 genes are indicated on the plots. B, Venn diagrams visualizing the overlap between R1881 vs CTRL (white) and R1881+RBN vs R1881 (grey) differentially expressed genes (p-adj < 0.001) and AR gene targets (blue) identified by incorporation of AR ChIP-seq and RNA-seq (R1881 treated, VCaP cells). The overall overlap is broken down into four sub-overlaps between specific combinations of genes with increased (red arrow) and decreased expression (blue arrow) genes in each group. C, The number of overlapping genes when differentially expressed genes between R1881 vs. CTRL and R1881+RBN vs. R1881 are compared. The overlap size on the y-axis is drawn as a step function over the respective ranks based on the Wald statistic. Top ranks correspond to genes with increased expression and bottom ranks to genes with decreased expression. Only the top 1,000 and the bottom 1,000 ranks are shown. In addition, the expected overlap and 95% confidence overlaps derived from a hypergeometric distribution are also shown. D, Heatmaps showing the top 30 genes that contribute to the similarity score in panel C (left - increased expression with R1881, right - decreased expression with R1881). From the left, columns represent CTRL, R1881, and R1881+RBN experimental groups. The values were normalized using the variance stabilizing transformation (VST) and presented. The representative color gradient used for the heatmap is presented in the figure.

The effects of PARP7 inhibition indicated a trend whereby expression of genes increased by R1881 was further increased by R1881+RBN2397, and similarly, genes which expression was decreased by R1881 was further decreased with RBN2397 co-treatment (Fig. 1B). The largest percentage of androgen-sensitive genes affected by the PARP7 inhibitor (45%) were those positively regulated by R1881 (Fig. 1B). To better understand the effects of PARP7 inhibition, we quantified the similarity between the ranked lists of genes from the two comparisons. Fig. 1C shows the number of shared top- and bottom-ranked genes based on the Wald statistic in the two comparisons, along with the expected overlap and 95% confidence intervals derived from a hypergeometric distribution. The number of shared genes with increased expression in this analysis is striking, suggesting that genes most elevated in expression by R1881 are also among the genes that are most elevated by PARP7 inhibition. Genes that contribute most to this similarity clearly exhibit a stronger RBN2397 effect compared to the response to R1881 alone in the same direction (Fig. 1D). We also used our previously generated RNA-seq from DTX3L knockdown (KD), R1881-treated VCaP cells, to test for enrichment in R1881+RBN2397 vs. R1881 DEGs (Supplementary Fig. 1B). We used DTX3L KD-sensitive genes from R1881-treated samples, excluding genes whose basal expression levels were affected by DTX3L KD, to create a gene set. We then performed gene set enrichment analysis (GSEA) to test the enrichment of R1881+RBN2397 vs. R1881 DEGs in that gene set. This analysis showed no enrichment. Taken together, these data suggest that PARP7 can act as a negative regulator of AR-dependent gene expression, and the effect is separable from the E3 function provided by DTX3L.

### Modules of AR-dependent gene expression

Our finding that genes whose expression was most affected by RBN2397 also showed a relatively small effect of R1881 alone in the same direction led us to hypothesize that PARP7 exerts negative feedback on AR, apparent at the 18-hr timepoint. PARP7 is known to exert negative feedback on AHR-regulated genes such as CYP1A1(*11*). We envisioned that negative feedback in prostate cancer cells, initiated by AR induction of PARP7, could temporally regulate the effects of circulating and tumor-derived androgens and potentially affect disease progression.

To study the temporal effects of androgen signaling and AR-dependent gene expression in VCaP cells, we used publicly available RNA-seq data (R1881 treatment for 8, 12, 18, 22, 24, 48 hr) to compile an androgen treatment time course (Fig. 2A). This enabled the investigation of AR-dependent gene expression on a transcriptome scale. We employed weighted gene co-expression network analysis (WGCNA) to identify modules of genes that co-express with similar time-dependence(*23*). The patterns are summarized as a weighted average expression profile termed a module eigengene, which is the first principal component of a given module. From the initial 32 modules rendered by the analysis, we retained 19 modules, each containing >100 genes (Supplementary Fig. 2).

**Fig. 2:**
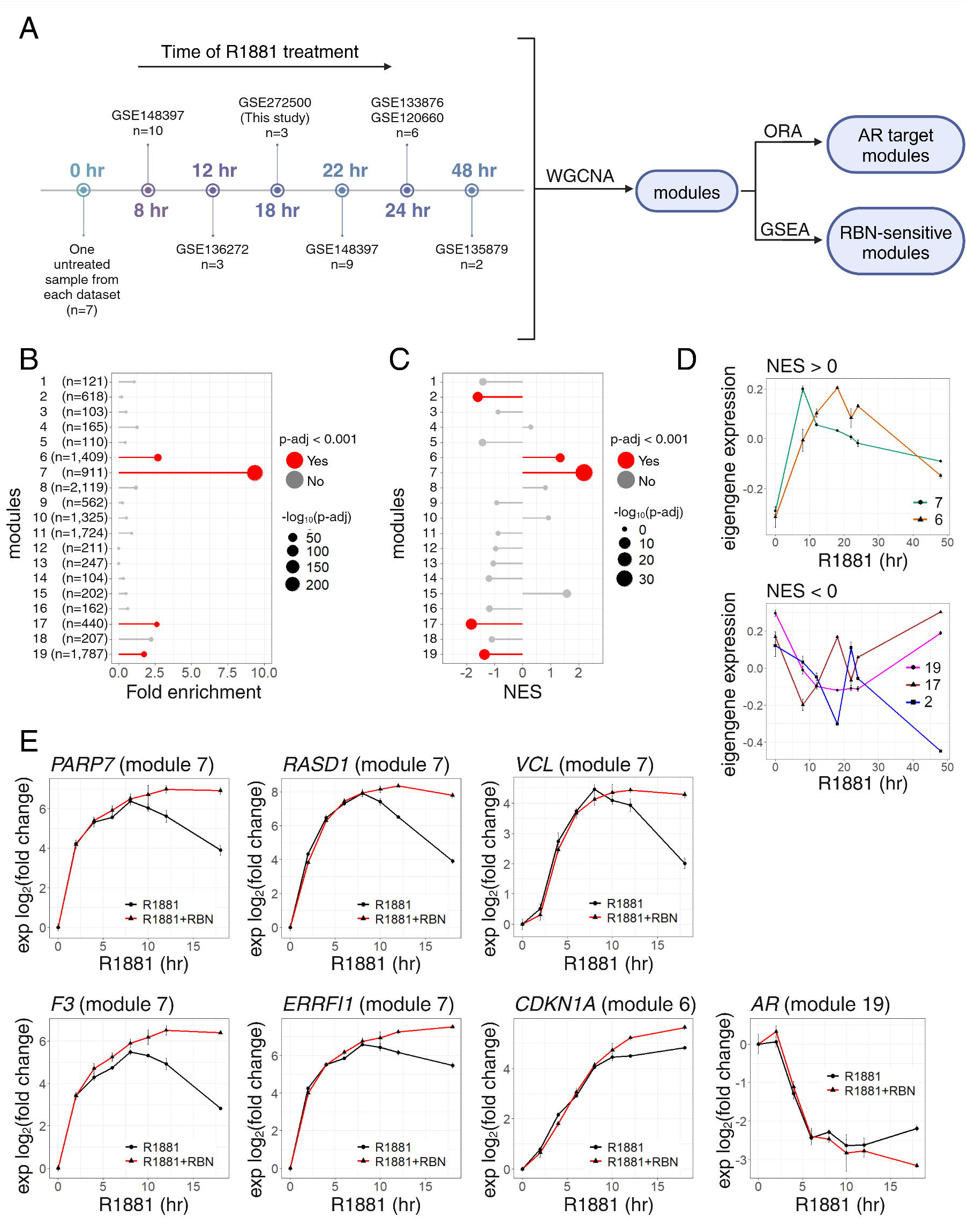
PARP7 regulates temporal patterns of AR-dependent gene expression through negative feedback. A, Scheme visualizing the analysis workflow. The left side presents RNA-seq datasets used to create the compiled gene expression time course of androgen treatment in the VCaP cell line. The GEO accession number and the number of samples are shown for each time point. The right side presents the analysis workflow (WGCNA – Weighted gene co-expression network analysis, ORA – Overrepresentation analysis, GSEA – Gene set enrichment analysis). B, Dot plot showing the gene modules overrepresented in the set of 1000 AR target genes. The gene lists of modules only with more than 100 genes resulting from WGCNA were used as gene sets for ORA. The y-axis represents the modules (the number (n) of genes is provided in parentheses for each module), and the x-axis represents the fold enrichment. The dot size represents -log10(p-adj), and the dots colored in red represent modules with p-adj < 0.001. C, Dot plot showing the gene modules enriched in the R1881+RBN vs R1881 differentially expressed genes. The gene lists of modules only with more than 100 genes resulting from WGCNA were used as gene sets for GSEA. The y-axis represents the modules, and the x-axis represents the Normalized enrichment score (NES). The dot size represents -log10(p-adj), and the dots colored in red represent modules with p-adj < 0.001. D, Line plot showing the eigengene expression for significantly enriched modules from GSEA in panel C. The plot on the top shows modules with positive NES (7 and 6), and the plot on the bottom shows modules with negative NES (19, 17, and 2). The y-axis represents the eigengene expression, and the x-axis represents the time of androgen treatment. Error bars represent standard deviation (0 hr: n = 7; 8 hr: n = 10; 12 hr: n = 3; 18 hr: n = 3; 22 hr: n = 9; 24 hr: n = 6; 48 hr: n = 2; n represents number of biological replicates). E, Line plots showing the results of a time course RT-qPCR experiment in VCaP cells treated with R1881 (black) and cotreated with R1881 and RBN3297 (red). The y-axis represents the log2 of the expression fold change from 0 hr time point normalized to the GUS housekeeping gene, and the x-axis represents the time of treatment in hours. The module membership is shown in parenthesis for each gene. Error bars represent standard deviation (n = 3; n represents number of biological replicates).

We first identified the modules that were enriched for AR targets. Using modules as gene sets, we tested for enrichment of the 1,000 AR target genes identified by CistromeGO in VCaP cells using over- representation analysis (ORA). Using a 0.001 p-adj cut-off, four modules (6, 7, 17, 19) emerged as significant, with Module 7 showing the highest fold-enrichment (Fig. 2B). Using GSEA, we also looked for modules that were enriched for R1881+RBN2397 vs. R1881 DEGs. Strikingly, this analysis identified the same modules and, additionally, module 2 (Fig. 2C). Plotting the eigengene expression values of the five modules as a function of R1881 treatment time showed that modules with positive enrichment (NES > 0) in RBN2397-sensitive genes showed a rapid increase in expression with peaks at 8 hr (module 7) and 18 hr (module 6), followed by a steady decrease in expression (Fig. 2D). In contrast, modules with negative enrichment (NES < 0) in RBN2397-sensitive genes showed reduced expression, with troughs at 8 hr (module 17) and 18 hr (modules 2, 19), and subsequent increase in expression. Overall, the five modules illustrate the kinetic effects of androgen signaling in VCaP cells, and integrating these findings with our RNA-seq data suggests that PARP7 activity has a prominent role in shaping the AR transcriptome.

### PARP7 activity and module kinetics

To validate the results from the compiled data set analysis, we performed a time course of R1881 and RBN2397 treatment in VCaP cells and used quantitative PCR (qPCR) to measure the expression of genes from select modules. We focused on a set of RBN2397-sensitive genes from module 7 since it had the highest enrichment score (Fig. 2B, C). We also included *CDKNA1* (module 6) and AR (module 19) for their biological and clinical relevance. All module 7 genes tested (*PARP7, RASD1, VCL, F3, ERRFI1*) displayed similar expression kinetics reflective of the module eigengene generated from compiled RNA-seq data sets (Fig. 2D, E). This included a steep induction, a peak of transcription at approximately 8 hr, and subsequent decrease, the effect of which was reduced by RBN2397 treatment (Fig. 2E). For comparison, the module 6 gene *CDKN1A* had peak expression at 18 hr with a slight fold-increase in response to RBN2397. Lastly, the module 19 gene AR showed a steep R1881-dependent drop and was only affected by RBN2397 during a late recovery phase (12-18 hr). These results confirm that PARP7 controls a negative feedback mechanism that acts preferentially on AR target genes.

### Androgen-induced degradation of AR

Given the temporal effects of androgen on certain modules of gene expression, we used immunoblotting to query if the transcriptional changes were associated with AR protein levels. In VCaP cells, we observed a decline in AR protein levels after ∼4 hr of R1881 treatment (Fig. 3A, D), consistent with prior work showing that AR represses its own transcription(*24*). The reduction in AR protein level was blunted by PARP7 inhibition (Fig. 3B, D). Though this finding suggested a link between PARP7 activity and AR degradation, ADP-ribosylated AR was not readily detected in extracts from VCaP cells exposed to R1881 in culture (Fig. 3A). This was despite the fact that AR undergoes androgen- and PARP7-dependent ADP-ribosylation in response to androgen(*12*). ADP-ribosylated AR was detected, however, when VCaP cells were co-treated with R1881 and the proteasome inhibitor Bortezomib (Fig. 3C, D; +R1881+Bortez). The splice variant AR-V7 in VCaP cells also showed an androgen-induced degradation and Bortez-dependent accumulation similar to full-length AR (Supplementary Fig. 3A-D). Additionally, AR levels were reduced in VCaP cells treated with nanomolar concentrations dihydrotestosterone (DHT), a physiological ligand that is more metabolically labile than the synthetic compound R1881 (Supplementary Fig. 3E).

**Fig. 3:**
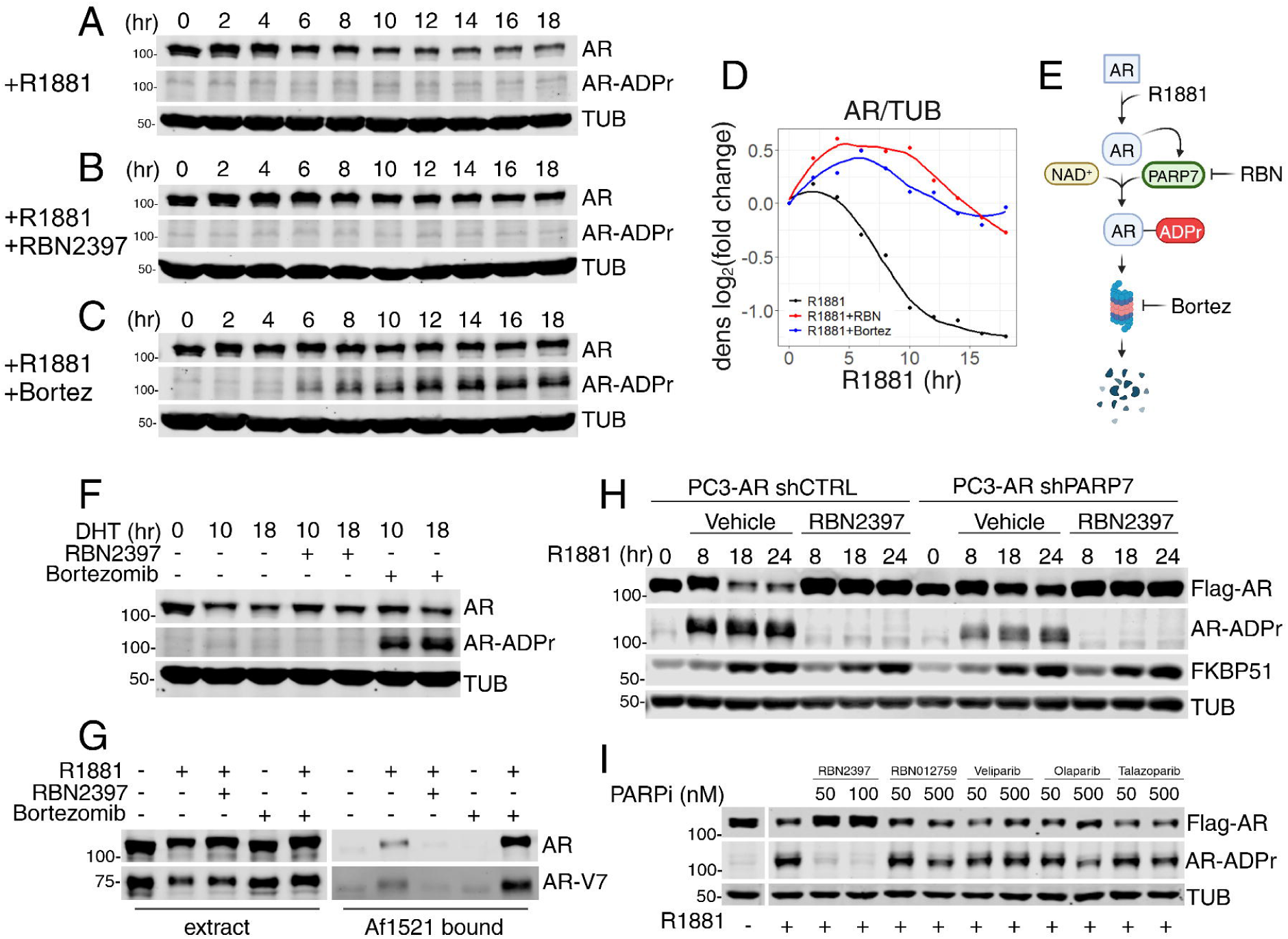
Androgen-induced ADP-ribosylation by PARP7 targets AR for proteasomal degradation. A-C, Immunoblot detection of AR and AR-ADPr (by FL-AF1521) in VCaP cells subjected to a time course of R1881 (A), R1881 and RBN2397 (B), or R1881 and Bortezomib (Bortez) (C) treatment. Cells were collected every 2 hours for an 18-hour period. D, Line plot visualizing the AR protein density measurements for immunoblots from panels A-C. The y-axis represents the log2 of the AR/TUB density fold change from 0-hr time point, and the x-axis represents the time of treatment in hours. The curves were fitted using the loess method. Data points represent single measurements. E, Schematic diagram of the androgen-induced AR degradation mechanism. F, Immunoblot detection of AR and AR-ADPr (Fl-AF1521) in VCaP cells treated with DHT, DHT+RBN2397, and DHT+Bortezomib for times indicated on the panel. G, Immunoblot detection of the AR and AR-V7 protein in VCaP cell extracts and AF1521 bound fractions. Cell extracts from VCaP cells treated with different combinations of R1881, RBN2397, and Bortezomib for 10 hr and were combined with AF1521 beads for the enrichment of ADP-ribosylated proteins. H, Immunoblot detection of Flag-AR, AR-ADPr , and FKBP51 in PC3-AR shCTRL (left) and shPARP7 (right) cells treated with R1881 and cotreated with R1881 and RBN2397 for times indicated on the panel. The shGFP was used as shCTRL. I, Immunoblot detection of Flag-AR and AR-ADPr in PC3-AR cells treated for 18 hr with R1881 and different PARP inhibitors (RBN2397 – PARP7i; RBN012759 – PARP14i; Veliparib, Olaparib and Talazoparib – PARP1/2i).

### PARP7 marks AR for destruction by the proteasome

DHT- and Bortez-co-treatment resulted in the accumulation of ADP-ribosylated AR, consistent with a model where ADP-ribosylated AR is marked for degradation by the proteasome (Fig. 3E, F). The model is supported by pull-down assays using the recombinant macrodomain from *Archaeoglobus fulgidus* immobilized on beads (GST-AF1521)(*25*). There was an increased recovery of ADP-ribosylated AR using extract from Bortez-treated cells (Fig. 3G). Partial depletion of PARP7 by shRNA(*21*) was sufficient to reduce the androgen enhancement of AR degradation; this was particularly evident at the 18 and 24 hr timepoints (Fig. 3H). Although LSD1 recruitment mediates transcriptional repression of the AR gene(*24*) the results with Bortez (Fig. 3C, E, F, G), together with the fact that androgen-induced reduction in AR protein level occurs with expression from an ectopic promoter, establishes an important contribution from a post-transcriptional, ADP-ribosylation mechanism. The absence of AR transcriptional repression in PC3- AR cells may explain why longer periods of androgen exposure are necessary for a reduction in AR protein levels (Supplementary Fig. 3F). Results with inhibitors to other PARP enzymes (PARP1, 2, 14), including several used clinically, support the conclusion that AR ADP-ribosylation and androgen-induced degradation can be attributed to PARP7 (Fig. 3I).

### Mathematical model of PARP7 regulation of AR

PARP7 targets AR for degradation and this appears to explain the reduced AR output, consistent with negative feedback, observed after 8 hr of androgen treatment. To explore whether our understanding of the PARP7-AR relationship can account for the timing of the negative feedback mechanism, we developed an ordinary differential equation (ODE) based computational model of AR-PARP7 interactions and downstream transcriptional outputs. The model architecture includes AR and PARP7, each undergoing degradation at a basal and faster rate(*26*). It was assumed that the cell contains 6.6 AR molecules per promoter (based on 20,000 AR molecules - all ligand-bound - that can associate with 3050 responsive promoters(*27*)). AR transcript levels (Fig. 2E) were included in the model since AR represses its own transcription(*24*). Other key assumptions were that PARP7 drives AR ADP-ribosylation and that AR- ADPr is degraded at a different rate from the total pool of AR. An androgen-induced agonist conformation and nuclear localization are required for AR ADP-ribosylation by PARP7, which in prostate cancer cells is primarily a nuclear enzyme(*14*). We hypothesized that the ADP-ribosylation reaction occurs on DNA since PARP7 contains a Cys3His zinc finger and can be trapped in a chromatin fraction when inhibited by RBN2397(*21*). However, compartmentalization of the reaction could have an important impact on the outcome of the model, and so we tested two different model architectures to determine whether PARP7 ADP-ribosylates AR prior to or after engagement with DNA.

Based on these assumptions, we compared two different models to assess their ability to recapitulate the experimental data (Fig. 4A). In Model 1, AR undergoes ADP-ribosylation by PARP7 while bound to promoters, while in Model 2, AR undergoes ADP-ribosylation by PARP7 in the nucleoplasm. In both models, we allow modified and unmodified forms of AR interact with promoters. The complete equations for the models are described in the Methods.

**Fig. 4:**
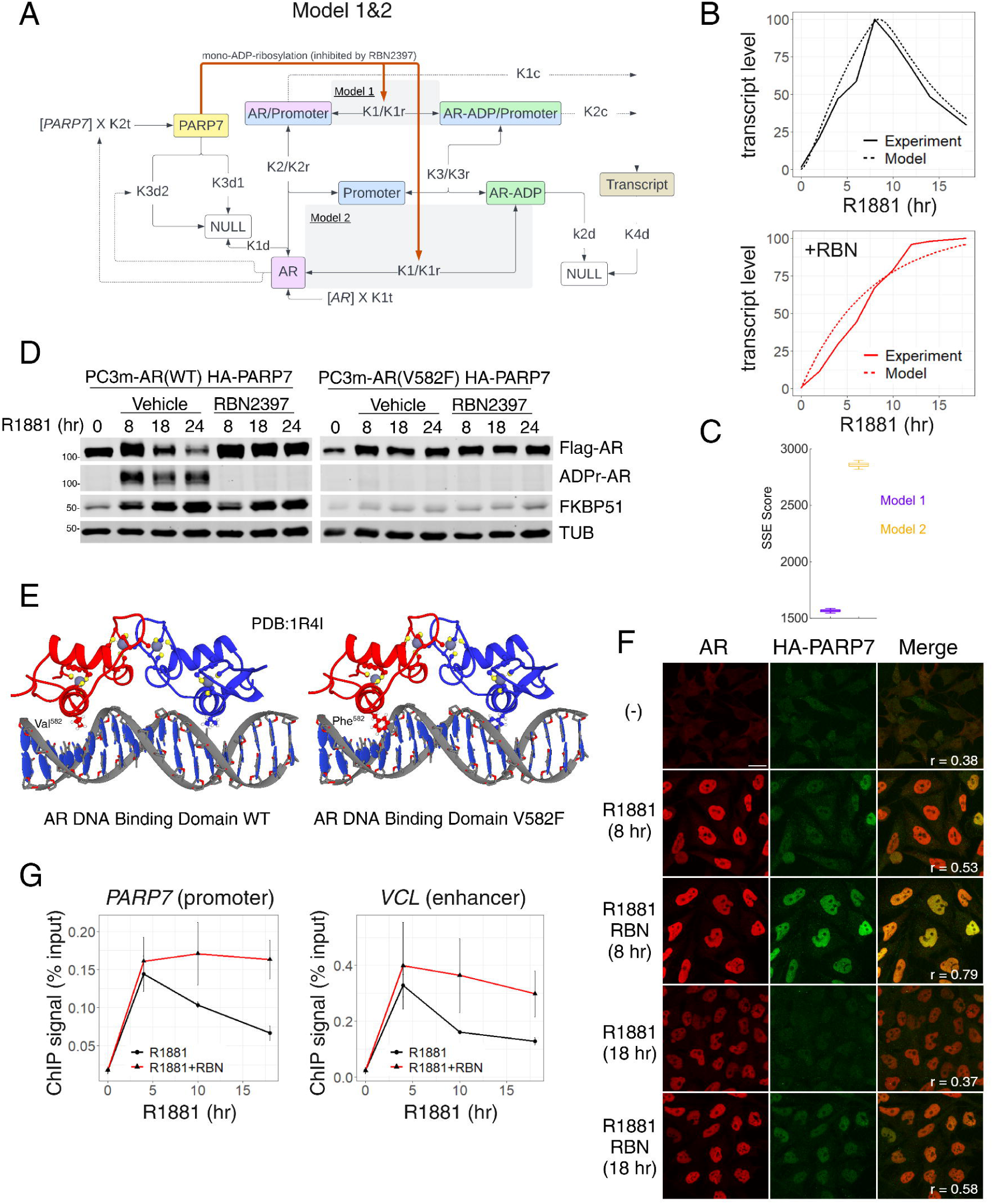
Androgen-induced AR degradation depends on DNA binding and affects chromatin occupancy. A, Schematic diagram of the model architecture that differs in the location where PARP7 can ADP-ribosylate AR, on promoter-containing chromatin in Model 1, or in the nucleoplasm in Model 2 (shaded boxes). Both modified and unmodified forms of AR can interact with promoters and generate transcripts. B, Plots of experimental transcript and simulated transcript during R1881 treatment, with and without RBN2397. Data is normalized to have a maximal value of 100. C, Graph of model performance for each model architecture, measured by the sum of squared error (SSE) after 600,000 Monte Carlo simulations to find the best parameters for each. The lower the SSE score, the better the simulated data matches the experimental data. For each model, top 100 scores were used to calculate p-vales (Wilcoxon test, n = 100; n represents repeated measurements). D, Immunoblot detection of Flag-AR, AR-ADPr (by FL-AF1521), and FKBP51 in PC3m(HA- PARP7/Flag-AR WT) cells (left) and PC3m(HA-PARP7/Flag-AR V582F mutant) cells (right) treated with R1881 or cotreated with R1881 and RBN2397 for times indicated on the panel. E, AR DNA binding domain structure (1R4I) in its wild-type form (left) and with V582F amino acid substitution (right). F, Confocal microscopy staining of HA-PARP7 and AR in PC3-AR(HA-PARP7) cells treated with R1881 or cotreated with R1881 and RBN2397 for times indicated on the panel. The third column shows merged channels, and the Pearson correlation coefficient for pixel colocalization is indicated in the bottom left corner for every condition. A scale bar of 10µm is provided on the bottom right corner of the upper left panel and applies to all panels. G, Line plots showing the results of a time course ChIP-qPCR experiment in VCaP cells treated with R1881 (black) or cotreated with R1881 and RBN3297 (red). The y-axis represents the ChIP signal characterized as the percentage of input DNA, and the x-axis represents the time of treatment in hours. Error bars represent standard deviation (n = 3; n represents number of biological replicates).

The parameters were fitted to each model using a Monte Carlo approach with 600,000 iterations (Fig. 4B; Supplementary Fig. 4A). Each model was able to give a qualitatively good fit to the transcriptional output of AR with and without RBN2397. Model 1, which restricts ADP-ribosylation to promoter-bound AR, provided the best fit to the experimental data, with the lowest sum of squares error (SSE) score (Fig. 4C). In both models, the parameters that best fit the data result in a scenario where ADP-ribosylated AR has a lower affinity for the promoter than unmodified AR. In model 1, the excess AR and a limited number of promoter binding sites provides a mechanism by which only a subset of AR molecules are available for PARP7 modification at a given time. Together with the low starting concentration of PARP7, this mechanism could explain the multi-hour delay of negative feedback, as seen in module 7. We compared the models by calculating their Bayesian Information Criterion (Supplementary Table 1), and then calculating the Bayes Weights to assess the best model while taking into account model complexity. The Bayes Weights were 0.95 for Model 1 and 0.05 for Model 2, indicating that the Model 1 architecture is more likely to be correct. Our mathematical modeling indicates that PARP7-induced degradation of AR is sufficient to capture the transcriptional effects of PARP7 inhibition given the listed assumptions about AR function, and suggests that ADP-ribosylation of AR likely occurs on chromatin.

### Validation that AR ADP-ribosylation requires DNA binding

We next tested the prediction from Model 1 that PARP7 ADP-ribosylation of AR depends on DNA binding, using an AR exon2 loss-of-function mutation (V582F) from a French family with complete androgen insensitivity syndrome that maps to the DNA recognition helix (Fig. 4D, E; described originally as V581F)(*28*). The V582F substitution prevents DNA binding, but it does not alter androgen affinity and androgen-induced AR phosphorylation(*29, 30*). In PC3m-AR(V582F) cells, the mutant AR showed virtually no androgen-induced ADP-ribosylation nor androgen-induced AR degradation (Fig. 4D). As expected, the DNA binding mutation abrogated androgen-induction of the AR target gene *FKBP51* (Fig. 4D).

Our data places ADP-ribosylation and the AR degradation mechanism in the context of chromatin. Accordingly, PARP7 activity is predicted to indirectly affect AR promoter occupancy by controlling androgen-induced AR degradation. We tested this by chromatin immunoprecipitation followed by qPCR (ChIP-qPCR) for AR as a function of androgen treatment time using two androgen-regulated genes that are RBN2397-sensitive (PARP7 and VCL). We observed that AR occupancy of these genes was maximal after ∼4 hr of androgen treatment, at which point the PARP7-mediated negative feedback became apparent (Fig. 4G). This effect was largely prevented by RBN2397 treatment (Fig. 4G). Thus, negative feedback via PARP7 manifests as a reduction in AR chromatin occupancy. Using confocal microscopy in a prostate cancer cell line expressing epitope-tagged AR and PARP7, we performed pixel-wise co-localization analysis. Based on Pearson’s correlation coefficients, AR and PARP7 distributions were correlated in cells treated with androgen (r=0.53) and increased when co-treated with RBN2397 (r=0.79)(Fig. 4F; Supplementary Fig. 4B). Taken together, these data suggest a mechanism for how androgen binding to AR in the cytoplasm culminates in AR ADP-ribosylation and degradation on chromatin.

### Site-specific cysteine ADP-ribosylation drives AR degradation

Previously we demonstrated that ADP-ribosyl-Cys modifications in the unstructured AR N-terminal domain (NTD) are recognized by macrodomains in the oligomeric DTX3L/PARP9 complex(*12*). To determine which ADP-ribosyl-Cys sites are responsible for androgen- and PARP7-mediated AR degradation, we used a panel of ten AR mutants with amino acid substitutions (Gly, Ser) for Cys sites shown by mass spectrometry(*12*) to undergo androgen-induced ADP-ribosylation in prostate cancer cells (Fig. 5A-F; Supplementary Fig. 5A-G, Supplementary Table 2). Glycine is inactive for ADP-ribosylation, while serine (similarity to cysteine) can serve as an ADP-ribose acceptor for some PARPs . Mutating eight ADP-ribosylation sites (Fig. 5B, F; Mut1) virtually eliminated androgen-induced AR ADP-ribosylation and degradation, but this set of mutations also reduced the transcription function of AR based on impaired induction of the direct target gene, *FKBP51* (Fig. 5A, B). Restoring amino acid 620 in Mut1 to Cys to create Mut2 rescued androgen-induced degradation and generated an ADP-ribose signal from AR detected by blotting using fluorescently labeled tandem AF1521 (FL-AF1521) (Fig. 5C, F) and by pull-down on AF1521 beads (Supplementary Fig. 5H). Substituting a Ser at position 620 in the context of a multi-site mutant (Mut3; Fig. 5D, F) or as a single-site mutant Cys620Ser (Mut4; Fig. 5E, F) failed to restore androgen-induced degradation, possibly because PARP7 shows a preference for Cys over Ser for ADP-ribosylation. Notably, mutation of Cys620 was sufficient to eliminate androgen-induced degradation of AR Mut10 (Supplementary Fig. 5F, G). This result demonstrates that Cys620 is required for androgen-induced degradation of AR, likely because it serves as an ADP-ribose acceptor site for PARP7.

**Fig. 5:**
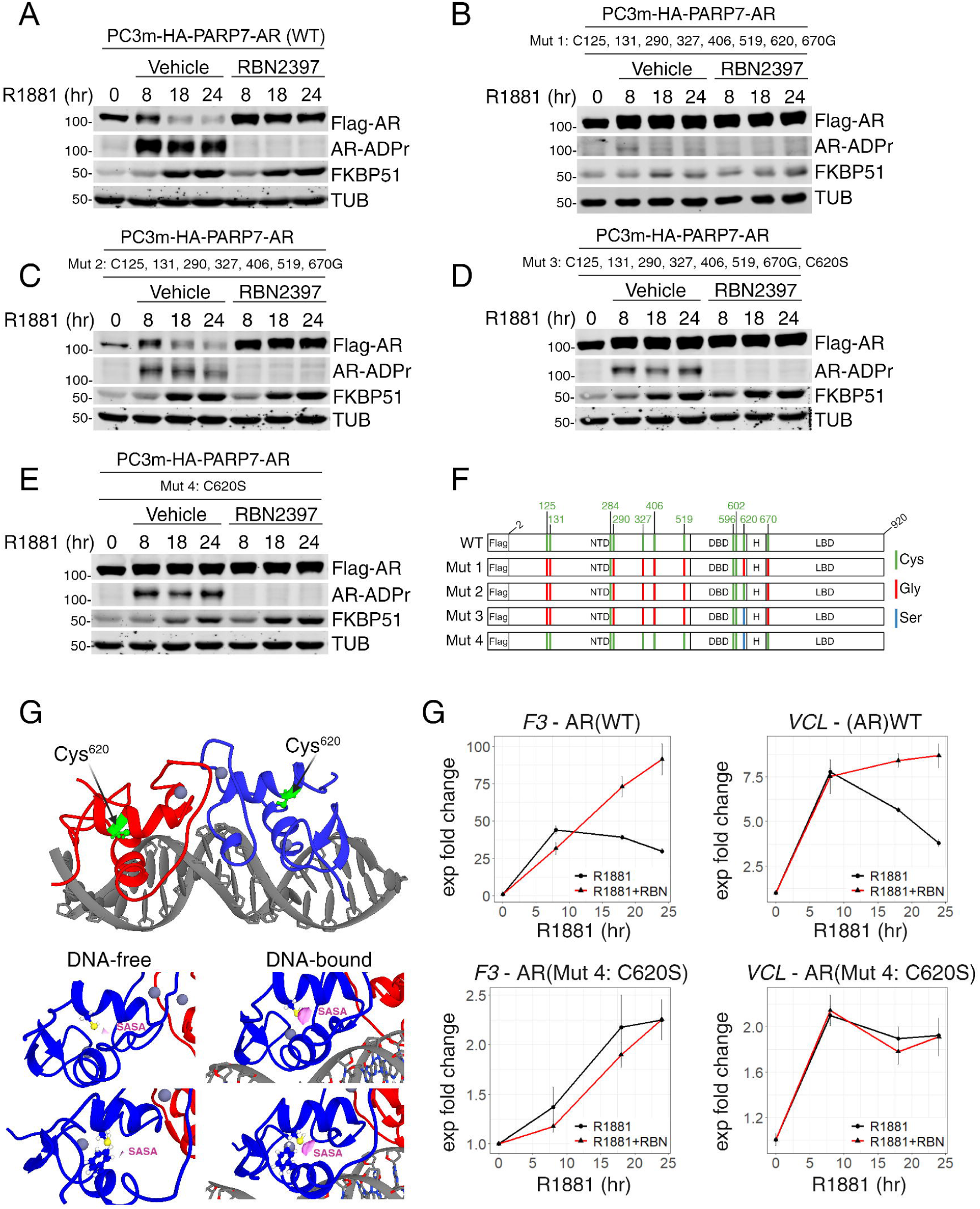
Mutation of the ADP-ribosylation site Cys620 eliminates androgen-induced AR degradation. A-E, Immunoblot detection of Flag-AR, AR-ADPr (by FL-AF1521), and FKBP51 in PC3m(HA- PARP7/Flag-AR wild type or mutant) cells treated with R1881 or cotreated with R1881 and RBN2397 for times indicated on the panel. A: AR WT; B: AR Mut1 (C125,131,290,327,406,519,620,670G); C: AR Mut2 (C125,131,290,327,406,519,670G); D: AR Mut3 (C125,131,290,327,406,519,670G & C620S); E: AR Mut4 (C620S). F, Diagrams of Flag-AR mutants employed in this figure (Mut 1-4). All of the ADP-ribosylation cysteine sites on AR are marked in green, and the particular substitutions are marked in red for glycine and blue for serine. G, Solvent accessible surface area (SASA) measurements of Cys620 in the co-crystal structure of the AR DNA binding domain (DBD) dimer. The upper panel shows the whole structure with Cys620 marked in green on each monomer. The middle panel shows the SASA (pink area) for the Cys620 when AR DBD is DNA-free (left) and DNA-bound (right). The panel at the bottom shows similar views but also displays the R group of Phe that is predicted to modulate the SASA of Cys620. H, Line plots showing the results of a time course RT-qPCR experiment in PC3m(HA-PARP7/Flag-AR WT) cells (upper panels) and PC3m(HA-PARP7/Flag-AR C620S mutant: Mut 4) cells (lower panels) treated with R1881 (black) and cotreated with R1881 and RBN2397 (red). The y-axis represents the log2 of the expression fold change from 0 hr time point normalized to the GUS housekeeping gene, and the x-axis represents the time of treatment in hours. Error bars represent standard deviation (n = 3; n represents number of biological replicates ).

### DNA binding increases the solvent accessibility of AR Cys620

A co-crystal structure of the AR DNA binding domain (DBD) bound to cognate DNA(*31*) showed that Cys620 localizes to an alpha-helix, though it is not a zinc coordinating residue and it does not contact DNA. The sulfur group in Cys620 can hydrogen bond with several atoms within the same alpha-helix, raising the question of whether DBD structure affects R-group accessibility for ADP-ribosylation by PARP7. ADP-ribosylated AR can be crosslinked and detected at the androgen-responsive promoter and enhancer sites(*12*), prompting us to consider whether DNA binding affects ADP-ribosylation site accessibility. We evaluated the solvent-accessible surface area (SASA) of Cys620 in the structure of the AR DBD bound to DNA, which was optimized with the GFN-FF force field(*32*) implemented in CREST(*33, 34*) (Fig. 5G, see Methods). We found that DNA binding increased the SASA of Cys620 from 0.0754 to 0.6631 Bohr^2^. There is a visible rotation of Phe584, which increases the SASA of Cys620.

DNA-based induction of ADP-ribosylation site accessibility to PARP7 could provide spatial regulation that limits the reaction to chromatin-bound AR.

While the gene module analysis was performed in VCaP cells, we determined that module 7 genes (*F3* and *VCL*) were similarly affected by PARP7 inhibition in the PC3 cell background (Fig. 5H, upper panels). In this setting, the Cys620 to Ser substitution stabilized AR (Fig. 5E) and rendered the androgen-dependent output of *F3* and *VCL* resistant to the effect of PARP7 inhibition (Fig. 5H). Taken together, our data link AR Cys620 to AR degradation, AR ADP-ribosylation, RBN2397 sensitivity, and gene output.

### E3 ligase DTX2 controls the cellular level of ADP-ribosylated AR

Multiple Ub E3 ligases can promote AR degradation,(*35–39*) though none require an androgen-induced conformation of AR for ubiquitylation. The dependence of AR degradation on AR ADP-ribosylation suggested that cells express an E3 which recognizes an ADP-ribosyl-based degron. To identify an E3 that selectively degrades ADP-ribosylated AR, we focused on E3’s that encode ADP-ribose-binding domains. The Deltex C-terminal domain (DTC) is an ADP-ribose binding module conserved in DELTEX family members (*40*), which is part of the RING-DTC that recruits ADP-ribosylated substrates for ubiquitylation(*16, 17*). We included three E3’s that encode the Trp-Trp-Glu (WWE) domain, which can bind poly-ADP-ribose(*41*). Also tested were two E3’s that lack ADP-ribose sensing domains but mediate degradation of AR(*35*) and other steroid hormone receptors(*42*). PC3-AR cells were transfected with siRNA to the E3’s (Supplementary Fig. 6A), treated overnight with androgen, and subsequently blotted for ADP-ribosylated and total AR. We predicted that the depletion of an E3 that selectively degrades ADP-ribosyl-AR would increase the AR-ADPr/AR ratio. Depletion of several E3’s resulted in AR-ADPr/AR ratios >1; however, DTX2 depletion generated an AR-ADPr/AR ratio >10 (Fig. 6A). A time course of androgen treatment in siCTRL and siDTX2 cells displayed a clear accumulation of ADP-ribosylated AR (Fig. 6B). We verified the finding by showing that DTX2 depletion increases the level of ADP-ribosylated AR recovered on AF1521 beads (Fig. 6C). To assess whether DTX2 uses its DTC domain to bind ADP-ribosylated AR, we made use of a DTC mutant that contains three amino acid substitutions(*17*) that eliminate ADP-ribose binding (Fig. 6E; Supplementary Fig. 6B). With a synthetic ADP-ribosylated AR peptide,(*43*) we used fluorescence polarization to confirm the DTC domain can bind ADP-ribosyl-Cys, that the interaction requires the ADP-ribose moiety, and that binding is lost in the DTC mutant (Supplementary Fig. 6C). WT and mutant GST-DTX2 RING-DTC proteins immobilized on GSH beads were used for pull-down assays with extracts from cells depleted of DTX2 and treated with androgen. In the pull-down assay, ADP-ribosylated AR binding to DTX2 depended on a functional DTC domain, and the level of binding was increased about 2-fold by DTX2 depletion (Fig. 6D). Because the ADP-ribose conjugated to AR by PARP7 is also recognized by the MAR antibody (Supplementary Fig. 6D), our data indicates the DTC domain of DTX2 recognizes mono-ADP-ribose conjugated to AR.

**Fig. 6:**
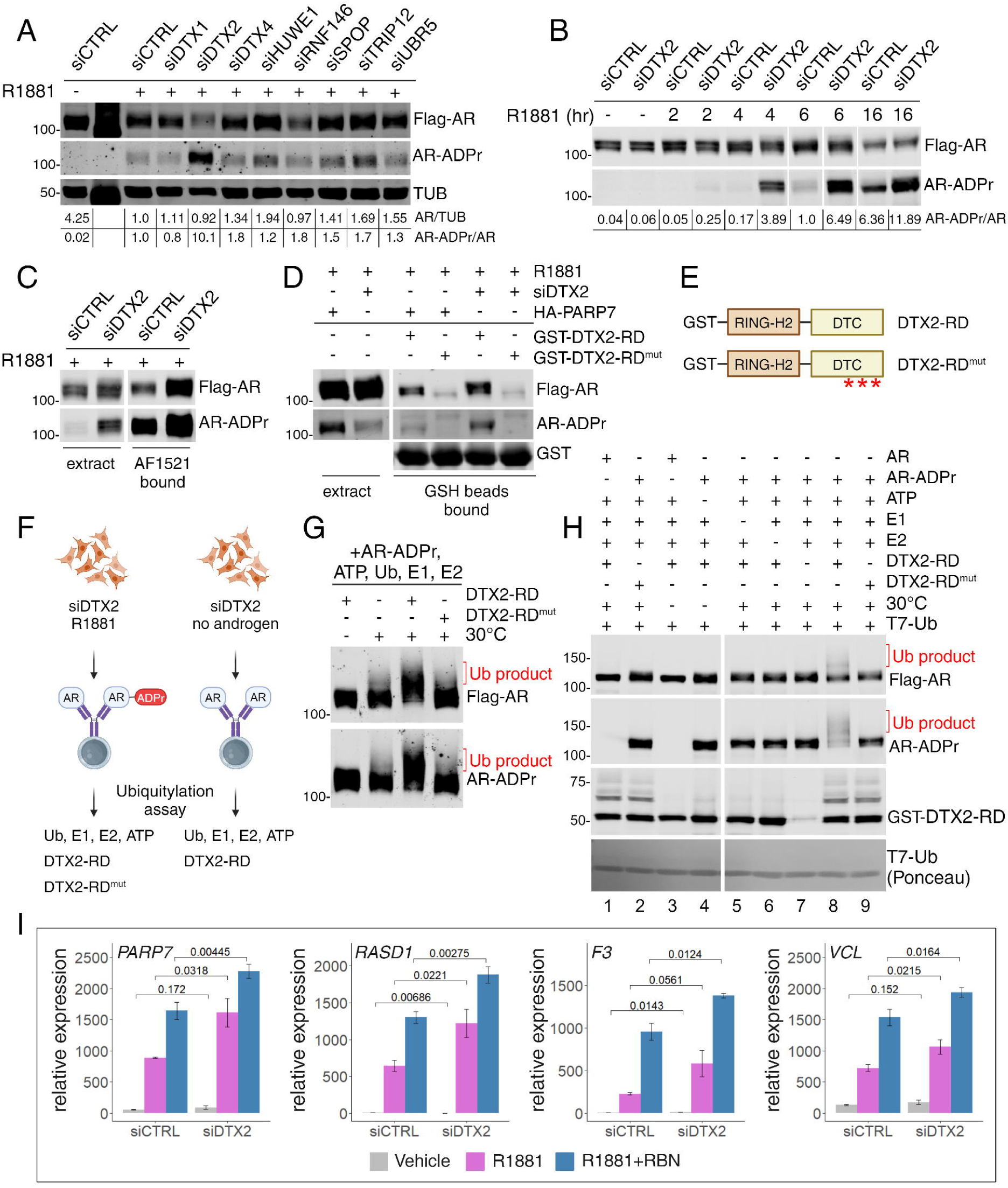
DTX2 is the E3 ligase for ADP-ribosylated AR. A, Immunoblot detection of Flag-AR and AR-ADPr (by FL-AF1521) in PC3-AR cells with siRNA knockdowns (total 4-day knockdown) of the selected relevant E3 ligases (DTX1, DTX2, DTX4, HUWE1, RNF146, SPOP, TRIP12 and UBR5), treated with R1881 for 21 hr before cell harvest. The AR/TUB and AR-ADPr/AR ratios for each lane are presented below the blot. B, Immunoblot detection of Flag-AR and AR-ADPr in PC3-AR cells with siDTX2 knockdown, treated with R1881 for times indicated on the panel. The AR-ADPr/AR ratio for each lane is presented below the blot. C, Immunoblot detection of Flag-AR and AR-ADPr in PC3-AR siCTRL and siDTX2 cell extracts and AF1521 bound fraction. Cell extracts from PC3-AR siCTRL and siDTX2 cells treated with R1881 for 6 hr were combined with AF1521 beads for the enrichment of ADP-ribosylated proteins. D, Immunoblot detection of Flag-AR and AR-ADPr in PC3-AR siCTRL/siDTX2 and PC3-AR HA- PARP7 cell extracts and GSH beads bound fraction. Cell extracts from PC3-AR siCTRL/siDTX2 and PC3-AR HA-PARP7 cells treated with R1881 for 6 hr were combined with GSH beads loaded with GST- DTX2-RD or GST-DTX2-RD^mut^ for the enrichment of proteins recognized by DTX2 DTC domain. E, Diagrams of DTX2-RD and DTX2-RD^mut^. Three loss of function mutations in the DTC domain of DTX2-RD^mut^ (S568A, H582A, and H594A) are indicated with red asterisk. F, Schematic diagram of AR protein preparation as a substrate for biochemical reactions. Cell extracts from PC3-AR siDTX2 cells treated (left) or untreated (right) with R1881 for 6 hr were combined with M2 beads for immunoprecipitation. The purified protein from the preparation with R1881 treatment was used for experiments in panels G and H, and the purified protein from the preparation without R1881 treatment was used only in panel H. G, Immunoblot detection of Flag-AR and AR-ADPr from the ubiquitylation assay on AR protein prepared with siDTX2 and R1881 treatment (panel F, left). The ubiquitylated products (Ub product, red bracket) are labeled for Flag-AR and AR-ADPr detection. All reactions contained AR-ADPr (R1881 treated samples), ATP, Ub, E1 and E2. For DTX2-RD status (dropout or DTX2-RD^mut^), refer to labels. H, Immunoblot detection of Flag-AR, AR-ADPr, and GST-DTX2-RD from the ubiquitylation assay on AR protein prepared with siDTX2 transfection, and with or without R1881 treatment (panel F). The ubiquitylated products (Ub product) are labeled in red for Flag-AR and AR-ADPr detection. The dropouts of the ubiquitylation assay components (Ub, E1, E2, and 30°C incubation) are indicated on the labels. The T7-Ubiquitin (T7-Ub) was detected by Ponceau staining. Lane numbers are indicated below the blot. I, Bar plots showing the results of an RT-qPCR experiment in PC3-AR siCTRL/siDTX2 cells untreated (grey), treated with R1881 (purple), and cotreated with R1881 and RBN2397 (blue). The y-axis represents the relative expression normalized to the GUS housekeeping gene, and the x-axis represents the siRNA used. The p-values from the Welch’s t-test for comparisons between corresponding conditions in siCTRL and siDTX2 are indicated on the plots. Error bars represent standard deviation (n = 3; n represents number of biological replicates).

### DTX2 uses its DTC and RING domains to ubiquitylate AR

To determine if DTX2 can conjugate Ub to ADP-ribosylated AR, we performed ubiquitylation assays with recombinant enzymes and AR immunoprecipitated (IP’d) from DTX2-depleted cells as the substrate (+R1881; Fig. 6F). When combined with Ub components, IP’d ADP-ribosylated AR underwent an upward gel shift indicative of ubiquitylation (Ub product) that was dependent on a functional DTC domain in DTX2 (Fig. 6G). Drop-out of individual Ub reaction components showed that AR undergoes androgen-, ADP-ribosyl-, and DTX2-dependent Ub modification which is dependent on a functional DTC domain (Fig. 6H; lanes 1, 2, 8, 9). These data suggest that DTX2 regulates AR levels and may contribute to transcriptional output by mediating the turnover of androgen-bound, ADP-ribosylated AR. We tested this prediction in PC3-AR cells by depleting DTX2 and assaying the effect on androgen-induction of Module 7 genes. By qPCR, we found that DTX2 depletion increased the androgen- and AR-dependent output of the *PARP7*, *RASD1*, *F3*, and *VCL* genes (Fig. 6I). The magnitude of the effects was less than observed with RBN2397 treatment, reflecting a partial depletion of DTX2 in these cells.

### ADP-ribosyl Cys is the acceptor site for ubiquitylation

Biochemical approaches were used to show that DTX family E3s can ubiquitylate substrates modified with ADP-ribose on serine(*16*) and acidic amino acids (*17*). To test if DTX2 can transfer Ub to ADP-ribosyl-Cys, we used a synthetic ADP-ribosylated AR peptide as a model substrate, which lacks lysine and a free amino terminus and cannot undergo canonical ubiquitylation (Fig. 7A, B). Ub conjugation to the AR peptide was found to be strictly dependent on the ADP-ribosyl-Cys moiety since no product was generated with an unmodified peptide or if the ADP-ribosyl peptide was pretreated with the phosphodiesterase NUDT16 which cleaves ADP-ribose (lanes 1-4; Fig. 7A, B; Supplementary Fig. 7A). Post-treatment with NUDT16 reduced the amount of Ub-peptide conjugate, consistent with Ub attachment via ADP-ribose (lanes 3, 5, 6; Fig. 7A). These data show that DTX2 conjugates Ub to ADP-ribosyl-Cys, and the substrate requires neither a lysine nor a free N-terminus.

**Fig. 7:**
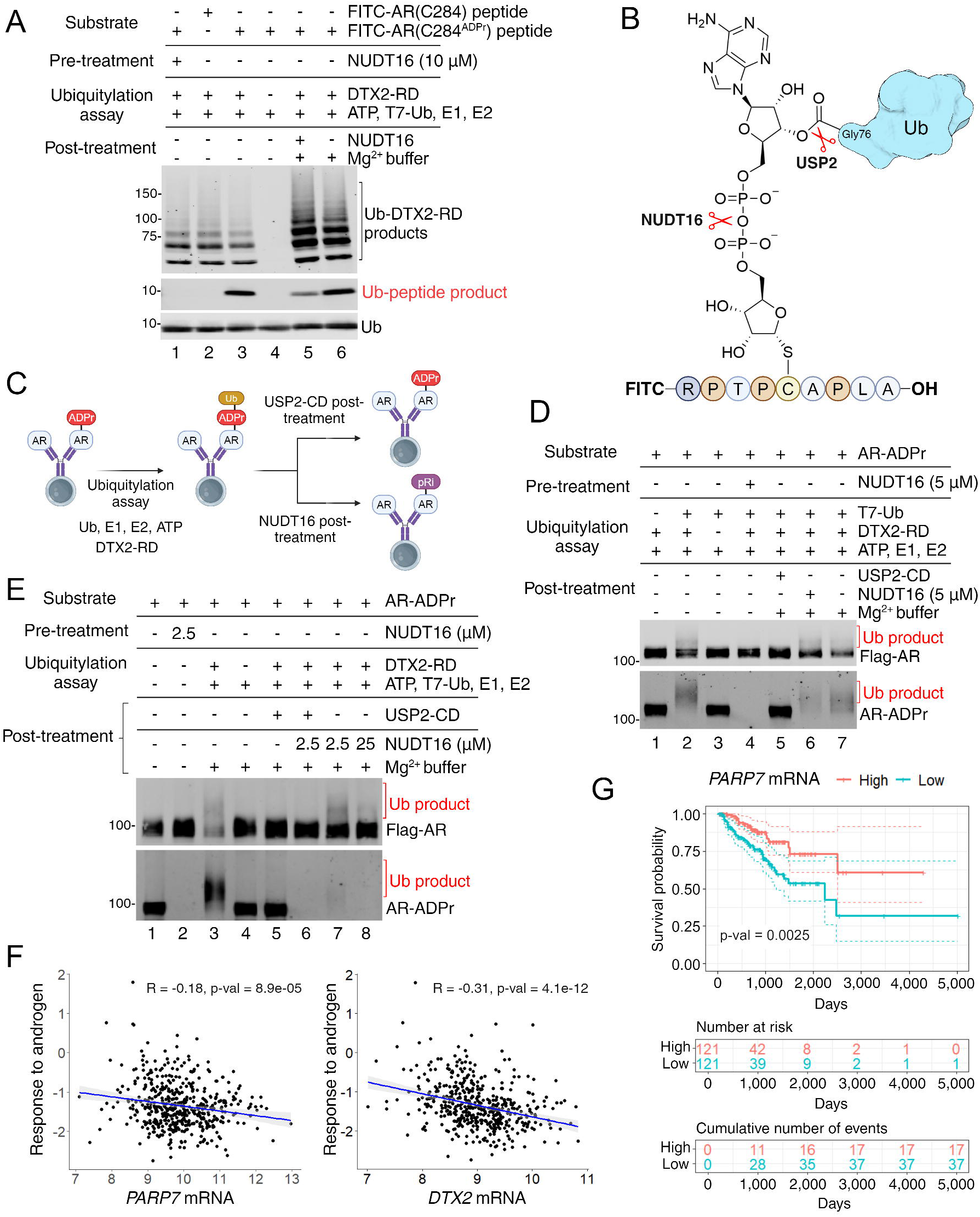
DTX2 conjugates ubiquitin to AR through ADP-ribose A, Immunoblot detection of Fluorescein (FITC) and T7-Ubiquitin (T7-Ub) from the ubiquitylation assay on FITC-AR(C284) or FITC-AR(C284^ADPr^) peptides. The labels indicate from the top: the substrate used (FITC-AR(C284) or FITC-AR(C284^ADPr^) peptides), pre-ubiquitylation assay treatments (NUDT16), the ubiquitylation assay (all reactions contained ATP, T7-Ub, E1 and E2, for DTX2-RD dropout, refer to labels), and the post- ubiquitylation assay treatments (NUDT16 or Mg^2+^ buffer). Lane numbers are indicated below the blot. B, Schematic diagram representing FITC-AR(C284^ADPr^) peptide conjugated to ubiquitin (Ub). Indicated with red scissors are bonds within the ADP-ribose structure cleaved by NUDT16 and USP2. C, Schematic of the ubiquitylation assay workflow for panels D and E. D, Immunoblot detection of Flag-AR and AR-ADPr (by FL-AF1521) from the ubiquitylation assay on AR protein prepared with siDTX2 transfection and R1881 treatment (refer to panel 6f for sample preparation workflow). The ubiquitylated products (Ub product) are labeled in red for Flag-AR and AR-ADPr detection. The labels separated by black lines indicate sequential steps from top to bottom. From the top, the labels indicate the substrate used (AR-ADPr), pre-ubiquitylation assay treatments (NUDT16), the ubiquitylation assay, and the post-ubiquitylation assay treatments (USP2-CD, NUDT16, or Mg^2+^ buffer). Lane numbers are indicated below the blot. E, Immunoblot detection of Flag-AR and AR-ADPr from the ubiquitylation assay on AR protein prepared with siDTX2 and R1881 treatment (refer to panel 6f for sample preparation workflow). The ubiquitylated products (Ub product) are labeled in red for Flag-AR and AR-ADPr detection. The labels separated by black lines indicate sequential steps from top to bottom. From the top, the labels indicate the substrate used (AR-ADPr), pre-ubiquitylation assay treatments (NUDT16), the ubiquitylation assay, and the post-ubiquitylation assay treatments (USP2-CD, NUDT16, or Mg^2+^ buffer). Lane numbers are indicated below the blot. F, Scatter plots depicting the correlation between PARP7 (left) or DTX2 (right) mRNA expression and the response to androgen pathway activity calculated by PARADIGM in primary prostate cancer patients from TCGA-PRAD cohort. Each dot represents one patient (n = 478, n represents number of patients). Pearson correlation coefficients and corresponding p-values are indicated on the plots. G, Kaplan-Meier plot depicting progression-free interval (PFI) in primary prostate cancer patients from the TCGA-PRAD cohort, stratified by PARP7 expression levels. The red line represents patients with high PARP7 expression (top 25%), and the green line represents patients with low PARP7 expression (bottom 25%). The X-axis represents time (days), and the Y-axis represents the progression-free interval probability. The interval distributions were compared using the log-rank test, with the p-value indicating statistical significance. Dotted lines represent the 95% confidence interval.

To further characterize the Ub linkage to ADP-ribosylated AR, we performed ubiquitylation assays with cell-derived ADP-ribosylated AR prepared by IP (see Fig. 6F). We included pre- and post-treatment with NUDT16, and post-treatment with the deubiquitylating enzyme USP2, which cleaves the Ub carboxyl terminus (Fig. 7B, C). A high molecular weight smear (Ub product) is detected when probing for AR and for ADPr; these products correspond to ubiquitylated AR since neither is detected with Ub drop-out (lanes 1, 2; Fig. 7D). The Ub product was eliminated by pretreatment with NUDT16 and by post-treatment with USP2 (lanes 4, 5; Fig. 7D). The Ub product displayed a slight resistance to post-treatment with NUDT16, as a smear was still detectable upon probing for AR and ADPr (lane 6; Fig. 7D). NUDT16 resistance could be indicative of Ub transfer to a proximal lysine (Supplementary Fig. 7B). The cleavage resistance of the Ub product was, however, overcome by increasing the NUDT16 concentration, temperature, and incubation time (lanes 3, 7, 8; Fig. 7E). These data are consistent with Ub conjugation to AR through an ADP-ribose linkage. Given how ADP-ribose sits in the binding pocket of NUDT16,(*44*) we speculate that Ub conjugated to the adenine proximal ribose might disfavor NUDT16 binding and partially protect the ADPr linkage from hydrolysis prior to engagement with DTX2. The proximal ribose ring in ADP-ribose is predicted to be the site of Ub attachment since, by modeling, binding to the DTC domain would place it in close proximity to the carboxyl terminus of Ub in the Ub-E2 complex.(*17*)

### PARP7 and DTX2 are negatively correlated with androgen signaling in prostate cancer

Our functional analyses in multiple preclinical models show that PARP7 and DTX2 are components of a negative feedback mechanism that helps limit androgen signaling through AR. To determine whether the expression of PARP7 and DTX2 affect androgen signaling in human prostate tumors, we focused on the data generated from prostate primary tumors in The Cancer Genome Atlas (TCGA-PRAD). We downloaded the batch effect-normalized mRNA data and the pathway activity scores generated using PARADIGM from UCSC Xena(*45, 46*) and queried the correlation between PARP7 and DTX2 mRNA expression and the “response to androgen” pathway activity. We found a weak negative correlation for TIPARP (R = -0.18) and a moderate negative correlation for DTX2 (R = -0.31) (Fig. 7F), showing that higher levels of these components are associated with a reduction in AR output. Using the log-rank test, we found significant differences in progression-free survival between patients stratified based on PARP7 gene expression in the TCGA-PRAD cohort (Fig. 7G). The effect of PARP7 expression on progression-free survival remains significant (HR = 0.56) even when we account for age of diagnosis in a multivariate Cox proportional hazard model. Our analysis shows that PARP7 expression is prognostic, and high TIPARP expression is associated with a better outcome (Fig. 7G). These conclusions fit our biochemical data and suggest PARP7 levels are correlated with better clinical outcomes through modulation of androgen signaling.

## DISCUSSION

We identified a post-transcriptional mechanism that regulates AR protein levels and gene expression in prostate cancer cells. The mechanism is based on androgen induction of PARP7, ADP-ribose writing by PARP7, ADP-ribose reading and ubiquitylation by DTX2, and degradation by the proteasome (Supplementary Fig. 7B). The mechanism results in post-transcriptional reduction of AR protein levels and is the basis of a PARP7-dependent negative feedback loop that controls the expression of specific modules of AR target genes. The data supporting these conclusions were derived from molecular, biochemical, and *in silico* approaches, which, taken together, provide new insight into how androgen signaling through AR can be regulated and generate temporal effects on gene expression. Prostate cancer cells therefore use three independent mechanisms to control AR levels: LSD1-mediated transcriptional repression of AR,(*24*) E3-mediated degradation of ligand-free AR(*35–39*), and ADP-ribosylation-dependent degradation of androgen-bound AR in association with chromatin. Importantly, the mechanism driven by androgen signaling limits degradation to AR that has fulfilled its transcription function.

One of the challenges for understanding PARP-dependent pathways is determining how family members locate and mediate substrate-specific ADP-ribosylation. Through mathematical modeling of the AR-PARP7 pathway and associated transcriptional outputs, we were able to infer that ADP-ribosylation occurs when AR is bound to DNA. We tested this hypothesis using a mutant form of AR that encodes an amino acid substitution in the DNA recognition helix (V582F) that results in loss-of-function and complete androgen insensitivity syndrome(*28*). Notably, the disease mutant AR binds androgen with nanomolar affinity, undergoes nuclear translocation, and conformation-dependent AF1 domain phosphorylation,(*29, 30*) but it fails to undergo ADP-ribosylation by PARP7. The AR V582F mutant, as a consequence, is not degraded in response to androgen treatment. The DNA binding requirement for AR ADP-ribosylation is also consistent with our observation that a subset of PARP7 is chromatin-associated and undergoes trapping in the nuclear fraction of cells treated with the catalytic inhibitor RBN2397(*20, 21*).

The ADP-ribosylation site that controls androgen-induced degradation (Cys620) is within the DNA binding domain. Mutation of Cys620 to serine eliminates androgen-induced AR degradation in cells. Although serine can act as an ADP-ribose acceptor with other PARPs, PARP7 shows a bias for ADP-ribosylation of Cys residues(*12, 47*). Using previously solved crystal structures of the AR DBD bound to DNA(*31*) and energy minimization approaches, we found that the solvent accessibility of Cys620 increases upon DNA binding. Restricting the formation of the ADP-ribosyl-Cys degron to chromatin- bound AR could couple the degradation mechanism to AR that has homo-dimerized on DNA and fulfilled its transcription function.

Although PARP1 is known to undergo multi-site auto-ADP-ribosylation, to our knowledge AR is the first substrate shown to undergo extensive, multi-site mono-ADP-ribosylation. Following the induction of the PARP7 gene, AR is ADP-ribosylated on a total of 11 sites, all of which are cysteines. Most of the ADP-ribosylation sites map to the unstructured NTD, which raises the question of how androgen binding to AR controls PARP7 engagement with sites that span a ∼400 amino acid unstructured region. Given the striking dependence of AR ADP-ribosylation on DNA binding, it appears that ADP-ribosylation of the NTD, like the DBD, occurs when AR is chromatin-bound. The fact that point mutations in the PARP7 zinc finger eliminate AR ADP-ribosylation(*14*) would be consistent with a chromatin targeting function for this domain. Chromatin might provide the context for PARP7 encounters with other transcription factors as well(*48, 49*), though this remains to be tested.

Our previous work, together with the current study, is beginning to shed light on the functional significance of multi-site ADP-ribosylation of AR, and in particular, the critical roles of reader proteins. ADP-ribosylated AR can be recognized by two different ADP-ribose readers. ADP-ribose on AR is read by the tandem macrodomains in the PARP9/DTX3L complex(*12*). Oligomeric assembly of PARP9/DTX3L, which would place at least six macrodomains in the complex, is required for efficient binding to ADP-ribosylated AR(*15*). In this study, we show the DTC domain of DTX2 also can read ADP-ribose conjugated to AR and that the reader function is obligatory for DTX2-dependent Ub conjugation to AR since these interactions are lost upon mutations in the ADP-ribose binding pocket of the DTC domain.

It is interesting to note that although DTX3L contains a DTC domain, prostate cancer cells lacking DTX3L still undergo androgen-induced AR degradation. This suggests that DTC domains, ADP-ribosylated substrates, or both, have features that impart selectivity for ADP-ribose reading. This consideration seems relevant to other reader-substrate interactions as well, given the large number of ADP-ribosylated proteins in the cell. PARP9/DTX3L binds to ADP-ribosylated AR through tandem macrodomains in PARP9, and in this setting, the DTC domain of DTX3L could be available to read and modify ADP-ribosyl-substrates in chromatin proximal to AR. Since DTX3L is required for p53 K48 Ub conjugation at DNA damage sites(*50*), it may have the capacity for both lysine-dependent and lysine-independent Ub conjugation, the latter utilizing a DTC domain.

Castrate-resistant prostate cancer (CRPC) remains a major clinical challenge. At the molecular level, CRPC generally relies on AR activity supported by intra-tumoral androgen synthesis, AR gene alterations to affect its sensitivity to ligands, and expression of therapy-resistant spliced forms such as AR-V7 that lack a ligand binding domain. Our work shows that in cells, PARP7, in effect, restrains AR function by initiating its degradation, and analysis of patient data suggests the mechanism may be relevant to clinical outcomes. Within the same cohort of patients, high PARP7 and high DTX2 expression in primary tumors are each negatively correlated with androgen signaling. Thus, high PARP7 could provide a survival benefit by limiting androgen signaling through a reduction of AR protein level. Interestingly, this is conceptually analogous to androgen deprivation therapy used clinically to reduce AR activity.

There is an emerging view that ADP-ribose writing, reading, and lysine-independent ubiquitylation are integrated as a protein degradation mechanism. Substrate engagement with a particular PARP enzyme and site-specific ADP-ribosylation act as substrate priming events that create an ADP-ribose degron. Ub conjugation then relies on selective recognition of ADP-ribose by an E3 ligase that contains a reader domain. In terms of the detailed mechanism, it was shown that DELTEX family E3 ligases conjugate the Gly76 carboxyl terminus of Ub to an oxygen atom in the adenine-proximal ribose(*17*). In the case of androgen-induced degradation of AR, mutation of a single ADP-ribosylation site (Cys620) protects AR protein from the aforementioned unconventional degradation. DTX2, the E3 ligase responsible for the degradation of ADP-ribosyl-AR, also senses and degrades DNA repair and chromatin-associated factors that, based on drug sensitivity, are ADP-ribosylated by PARP1(*16*). ADP-ribosyl-dependent degradation is expanding the conceptual framework for cellular functions mediated by the PARP family, and it raises the question of whether PARP inhibitor effects on protein homeostasis might evolve as a therapeutic consideration. Understanding ADP-ribosyl-dependent degradation mechanisms could complement the current rationale for using PARP1/2 inhibitors that are based on synthetic lethality associated with defective DNA repair.

## MATERIALS AND METHODS

### Cell culture

HEK293T (ATCC CRL-3216) was maintained in DMEM, 5% fetal bovine serum (FBS), and 1% Pen/Strep (P/S). PC3 (ATCC CRL-1435), PC3M (CVCL_9555), and derivative cell lines were grown in RPMI1640, 5% FBS, and 1% P/S. VCaP (ATCC CRL-2876) was grown in DMEM/F12, 10% FBS, and 1% P/S. All cells were grown in the presence of 5% CO2 and at 37°C. For RNA-seq and qPCR experiments, cell growth media were switched to phenol red-free medium and FBS depleted of androgens 24 h before the drug treatments. HEK293T are female and all other cell lines used in this study are male.

### Stable cell line generation

A 60 mm-plate of HEK293T cells (60-70% confluency) were transfected with 1.5 g of a lentiviral plasmid DNA carrying target gene and 0.75 g each of two accessory plasmids pMD2g and psPAX2 using ViaFect transfection reagent (Promega E498A, total DNA : ViaFect ratio = 1 g : 3 l) and fresh medium for 16 hrs, replenished with a high FBS growth medium (DMEM, 35% fetal bovine serum and 1% P/S) and grow for another 24 hrs. The lentivirus-containing medium was moved to a 10 ml-conical tube which was centrifuged (700 x g) to remove the cell debris. The supernatant was passed through a 0.45 m-EZFlow Syringe Filter, concentrated using Lenti-X Concentrator (Takara, 631232) and then transfected into a target cell line with 8 g/ml of polybrene in the growth medium for 24 hrs. After cells grow in the growth medium for 2-3 duplication time, cells were started selection with 1-2 g/ml of puromycin or 0.2 mg/ml of hygromycin in the growth medium according to its respective antibiotic selection marker on the plasmid DNA. For shRNA knockdown cell lines following shRNAs were used: shGFP (shCTRL):CCGGTACAACAGCCACAACGTCTATCTCGAGATAGACGTTGTGGCTGTTGTATTTTT, and shPARP7:CCGGGAAGGCAAGCTACTCTCATAACTCGAGTTATGAGAGTAGCTTGCCTTCTTTTT.

### siRNA transfection

Cells were transfected with 20 nM of siRNA using Lipofectamin RNAiMAX transfection reagent (Invitrogen 13778-075) and passaged when they reach confluency. Cells were grown for 4 days in total, and drug-treatments were done before cell harvest.

### RNA-seq

For RNA-sequencing, VCaP cells were plated in the phenol-free DMEM media supplemented with 10% charcoal-stripped (depleted of androgens) FBS and treated with R1881, RBN2397, and R1881+RBN2397 and corresponding vehicles (ethanol for R1881 and DMSO for RBN2397) for 18 hr in biological triplicates (distinct samples). RNA was extracted using a Qiagen RNeasy kit. Library preparation and sequencing were performed by Hudson Alpha. In brief, a fluorometric assay was used to assess the RNA concentration and integrity; the standard polyA method was used to make indexed libraries; size and concentration were determined using quality control, and samples were sequenced using Illumina HiSeq 2500 at a depth of 250 million × 50-bp paired-end reads. Reads were pseudoaligned to the hg38 genome (ENSEMBL GRCh38.89) using kallisto (*51*). The transcript-level abundance estimates were summarized to gene-level counts using tximport R package (*52*). The DESeq2 R package(*53*) was used to apply Variance-stabilized transformation (VST) to counts data and to determine differentially expressed genes between conditions. Genes with less than 20 counts in all conditions were eliminated (pre-filtering). For all downstream analyses, differentially expressed genes (DEGs) were additionally filtered (baseMean > 100). Volcano plots were generated using the EnhancedVolcano R package.(*54*) Venn diagrams were generated using eulerr R package(*55*) on filtered genes (p-adj < 0.001). All heatmaps were generated using the pheatmap R package(*56*) with VST counts.

### AR targets

Cistrome-GO (http://go.cistrome.org/) online tool was used to create a set of 1,000 AR target genes, allowing the integration of ChIP-seq and RNA-seq data. A VCaP ChIP-seq peak bed file was taken from a publicly available dataset (GSE84432, 4 hr of R1881 treatment) and the differential expression file (DESeq2) was also generated from a publicly available dataset (GSE148397, 8 hr of R1881 treatment).

Analysis was conducted with a half-decay distance of 10kb. The top 1,000 genes from the resulting analysis were used to create a list of AR targets.

### Pathway enrichment

All gene set enrichment analyses (GSEA) were performed using fgsea package(*57*) with nPermSimple = 500,000 and log2FoldChange as a gene-level statistic. Gene sets for GSEA in Fig. 1c were created by filtering the R1881 vs. CTRL DEGs (p-adj < 0.001, log2FoldChange > 0 for increased gene expression, log2FoldChange < 0 for decreased gene expression). To create a gene set for Supplementary Fig. 1B, our previously published RNA-seq dataset from VCaP cells +/- R1881 and +/- DTX3L knockdown (KD) (GSE133876) was used. DTX3L KD vs. CTRL DEGs were eliminated from R1881+DTX3L KD vs. R1881 DEGs (baseMean > 100, p-adj < 0.001) to enrich specifically for DTX3L-regulated genes that are induced by R1881. The gene set was created from the gene names of the remaining DEGs. Over- representation analysis (ORA) was performed using fgsea R package. The expected frequency was calculated by dividing the gene set (module) size by the background set size (24,824 genes) and multiplied by the study set (AR targets) size ((module size/24,824) x 1,000). To calculate the fold enrichment, the overlap between each module and AR targets was divided by the expected frequency.

### Compiling RNA-seq datasets

To create the compiled RNA-seq androgen treatment time course, the following publicly available RNA-seq datasets from VCaP cells were used: 8 hr - GSE148397 (n=10), 12 hr - GSE136272 (n=3), 18 hr – GSE272500 (this study, n=3), 22 hr- GSE148397 (n=9), 24 hr - GSE133876 (n=3) and GSE120660 (n=3) and 48 hr - GSE135879 (n=2). One untreated control from each study (n=7) was used as 0-hr. All datasets were uniformly reprocessed. FASTQ reads were pseudoaligned to the hg38 genome (ENSEMBL GRCh38.89) using kallisto. The transcript-level abundance estimates were summarized to gene-level counts using tximport R package. The resulting counts matrix was next batch-corrected using the ComBat R package.(*58*)

### Weighted gene co-expression network analyses

For weighted gene co-expression network analyses (WGCNA), the WGCNA R package(*23*) was used. The compiled counts matrix was pre-filtered to keep genes with 15 or more counts in at least 75% of all samples (30) or 75 or more counts in at least one sample. Next, the VST was applied to normalize the data. The soft power (β value) of 14 was selected based on the appearance of diagnostic plots (scale independence and mean connectivity). The blockwiseModules function was used to create the signed network, with Pearson correlation as a measure of similarity between genes, and to identify modules using hierarchical clustering on a signed topological overlap matrix. The max block size was set to 14,000 to process all the genes (13,811) in one block. Similar modules were merged based on their module eigengene, using a dendrogram cut height of 0.25 resulting in 32 modules. Modules with less than 100 genes were excluded, leaving 19 modules, from which a GMT file for GSEA and ORA was created.

### Tumor analysis

The TCGA-PRAD (The Cancer Genome Atlas – Prostate Adenocarcinoma) cohort was used for the patient data analysis. Log2 transformed expression data, PARADIGM pathway activity, and survival data were downloaded from Xena.(*46*) Scatter plots and Pearson correlation were generated with ggpubr R package.(*59*) The multivariate Cox model was built with progression-free interval (PFI) data using the survival R package(*60*). The Kaplan-Meier (KM) plot was generated with the survminer R package(*61*).

### Mathematical Modeling

Assumptions made: 1) All AR is considered to be ligand-bound. 2) AR degradation in the absence of ADP ribosylation is so slow as to be disregarded. 3) PARP7 has two degradation rates, a fast and slow rate. AR promotes the slow rate. 4) For Model 1, AR can only be ADP-ribosylated when bound to a promoter. 5) For Model 2, AR is ADP-ribosylated when not bound to a promoter. Pre-processing experimental datasets: Simulations were scored on their ability to predict the dynamics of the AR responsive transcripts PARP7, F3, and ERRFI1. Since PARP7, F3, and ERRFI transcripts have similar expression levels throughout the experiments, we averaged them and used that as a composite transcriptional profile for the model (referred to as PA7F3ER1 in the model). Additional time points in the transcriptional response were interpolated to score the simulations. The experimental datasets were normalized to have a maximum value of 100.

Simulations: Modeling was performed in MATLAB (version R2023b) using the ODE solver ode23s to solve the ODE system of mass-action equations (see Supplementary Table 1). The ODE solver runs 2,200 times with 0 to 0.01 integration, and each 100 runs represents an hour worth of transcript expression. A second model was built to capture the PARP7 inhibition by RBN2397. The two models run simultaneously sharing the same parameters and initial concentrations, but with the rate constant for ADP-ribosylation set to 0 for the inhibitor model. The two models output separate SSE scores which are then summed to assess the ability of the model to fit both + and – drug conditions. Initial Concentrations: All transcripts/proteins were initiated with zeros except AR and promoters. The number of AR responsive promoters in a cell has been found to be 3,050 and the number of AR molecules in the cells has been found to be 20,000, which leads to a ratio of 6.6 AR per promoter (*27, 62*). We maintained this ratio and found that a condition of 660 AR and 100 promoters was effective for performing the simulations.

Parameter Estimation: We divided the parameter estimation into two steps; initial parameters were determined by Monte Carlo selecting from a uniformly distributed parameters, followed by refinement using a Monte Carlo Markov Chain selecting from normally distributed parameters (*63*). We first ran 100,000 Monte Carlo simulations. The parameter sets were ranked from the lowest to highest SSE scoring, and we then chose the top 100 parameter sets. The last step is Monte Carlo Markov Chain method, more specifically Metropolis-Hastings algorithm (*64*). Each parameter set runs 5,000 times to explore its close surroundings and find the optimum parameter set. To ensure each worker does not generate overlapping random numbers, we initialized rng(), a random number generator, with a unique seed. The seed was calculated by multiplying the current second, using clock built-in function, by 1,000 and rounding down to an integer. We add the number of iterations to the seed to ensure different seeds if workers are initiated simultaneously. The top scoring parameter set for each model are shown in Supplementary Table 1. To select the best model, we calculated the Bayesian Information Criterion (BIC)(*65*). BIC is used as a method to assess the models using a maximum likelihood function.(*66*) The model with the lowest BIC is more likely correct. Bayes weights were calculated to provide a probabilistic measure (a normalized score between 0 and 1) of the competing models. The highest Bayes weight indicates the best model among the considered models. BIC and Bayes weights for each model are shown in Supplementary Table 1.

### Protein modeling

We used the crystal structure of the homodimer of the AR DNA binding domain (AR DBD) complexed with a DNA response element (DRE) from the Protein Data Bank (PDB ID: 1R4I). As the PDB structure originates from Rattus norvegicus and our experimental system pertains to Homo sapiens, there is a numbering shift of 18 amino acids. Additionally, we introduced an amino acid substitution in both protein chains, with alanine (A) for cysteine (C) at position 552 (PDB ID: 1R4I), aligning it with the structure of Homo sapiens. Subsequently, we optimized these structures following each substitution: A552C, A552C + C602S, A552C + V564F (numbering from PDBID: 1R4I) with the GFN-FF(*32*) force field implemented in the CREST program(*33, 34*) with the Analytical Linearized Poisson-Boltzmann (ALPB) implicit solvation modeling(*67*). The charge equilibrium model (EEQ) implemented in GFN-FF has difficulty distributing high molecular charge adequately over a large system. Hence, charges were initialized from the more robust charge extended Hückel method(*68*), originally developed in the context of the charge- adaptive q-vSZP basis set. Finally, we prepared six structures: three mutants each with and without DNA. We further calculated solvent accessible surface area (SASA) with and without DNA.

### RT-qPCR

Cells were harvested, the lysates were homogenized using a QIAshredder kit (Qiagen, 79656) and the genomic DNA was digested with DNase (Roche, 04716728001) according to the manufacturer protocol. The RNA was purified using an RNeasy kit (Qiagen, 74104). The cDNA synthesis was performed using an iScript cDNA synthesis kit (Bio-Rad, 1708890). The amplification of cDNA was performed using pairs of primers for *PARP7* (Forward; 5’- TCTCAGGAGCACTTGGAAAGA-3’, Reverse; 5’- TCAGCCTTCGTAGTTGGTCA-3’), *RASD1* (Forward; 5’-CCGCAAGTTCTACTCCATCC-3’, Reverse; 5’-TGAACACCAGGATGAAAACG-3’), *VCL* (Forward; 5’-TGACATTCTACGTTCCCTTGG-3’, Reverse; 5’-TTGGTTTTGGTCTGCAGGTT-3’), *F3* (Forward; 5’-GAACCCAAACCCGTCAATC-3’, Reverse; 5’-CGTCTGCTTCACATCCTTCA-3’), *ERRFI1* (Forward; 5’-GGAGCGCCTAATACCACTTG- 3’, Reverse; 5’-CCATTCATCGGAGCAGATTT-3’), *CDKN1A* (Forward; 5’- CTGCCGAAGTCAGTTCCTTG-3’, Reverse; 5’-CATGGGTTCTGACGGACAT-3’), and *AR* (Forward; 5’-GCCTTGCTCTCTAGCCTCAA-3’, Reverse; 5’-TGAATGACAGCCATCTGGTC-3’) in the presence of the HotStart™ 2X Green qPCR Master Mix (APExBIO, K1070), all according to manufacturer instructions. Technical replicates were averaged and normalized to housekeeping genes GUS (Forward; 5’-CCGACTTCTCTGACAACCGACG-3’, Reverse; 5’-AGCCGACAAAATGCCGCAGACG-3’).

Calculations were done using the comparative CT method. Reported are either fold change or relative expression values, depending which were more appropriate for the experimental question. Where needed, fold changes were log2 transformed to ensure a continuous scale for genes with increased and decreased expression. In each experiment, three biological replicates were used for each condition.

### Chromatin immunoprecipitation (ChIP)

VCaP cells were fixed using 1% formaldehyde in a growth medium at 37 °C for 10 min and neutralized with 125 mM glycine at room temperature for 5 min. Cells were harvested in cold PBS + 1 mM PMSF at 4 °C by centrifugation (2,200 × g, 4 °C, 5 min), resuspended in the extraction buffer (20 mM Tris pH 7.5, 1 mM EDTA, 100 mM NaCl, 2 mM DTT, 0.5% Triton X-100, and protease inhibitors) with end-over-end rotation at 4 °C for 20 min. Pellets were collected by centrifugation (2,200 × g, 4 °C, 5 min), washed with EDTA-free extraction buffer to remove the EDTA, and then sonicated (0.4-ml volume, Scale 4, 50% cycle, 10 pulse; repeat two more times). Samples were supplemented with CaCl2 (4 mM final) and micrococcal nuclease (NEB 500 gel units) to cleave the DNA at 37 °C for 5 min. EGTA (50 mM final) and SDS (0.1%) were added to quench the nuclease reaction. The samples were incubated for 10 min at 4 °C, followed by centrifugation (16,800 × g, 4 °C, 20 min). The supernatant was pre-absorbed with unconjugated Protein G beads in the presence of BSA (1 mg/ml final) and salmon sperm DNA (0.2 mg/ml final), then subjected to antibody bead binding at 4 °C for overnight. Beads were collected with centrifugation, washed five times with the extraction buffer supplemented with 0.1% SDS, then resuspended in 100 µl of reverse cross- linking buffer (125 mM Tris pH 6.8, 5% BME, and 1% SDS), and incubated at 65 °C for 6 hr. DNA was purified using the QIAquick Gel Extraction kit (Qiagen, 28704). The amplification of DNA was performed using pairs of primers for the *PARP7* promoter region (Forward; 5’- ACAAGGCCCACGAAATAGTC-3’, Reverse; 5’-CACCCTGTGAGGAAGCAAAC-3’), and *VCL* enhancer region (Forward; 5’-TGTGAGTTGGTGCTGCATAC-3’, Reverse; 5’- GGGAGTCAGGAACACAGAGT-3’) in the presence of the HotStart™ 2X Green qPCR Master Mix (APExBIO, K1070), all according to manufacturer instructions. Primers were designed based on AR ChIP-seq binding profiles in VCaP cells from a publicly available dataset (GSE84432, 4 hr of R1881 treatment). In each experiment, for every condition there were three biological replicates (distinct samples).

### Western blotting and AR-ADPr detection

After drug treatments, the media was removed, and the cells were lysed using a 1× SDS loading buffer (200 mmol/L Tris-HCl pH 6.8, 2% SDS, 10% glycerol, and 3% β-mercaptoethanol). The lysates were then heated at 95°C for 5 minutes, sonicated, centrifuged, separated by SDS-PAGE, and transferred to a nitrocellulose membrane. The membrane was blocked in PBS with 0.15% Tween-20 (PBST) and 5% nonfat milk at room temperature for 45 minutes. For standard western blotting, it was then incubated with 1° Ab/1% BSA/PBST for either 2 hr at room temperature or overnight at 4°C, washed at room temperature with PBST (five washes of 5 minutes each), then incubated with fluorescently labeled 2° Ab/1% nonfat milk /PBST for one hour at room temperature, and washed at room temperature with PBST (three washes of 5 minutes each). For AR-ADPr detection, it was incubated overnight at 4°C with 1 μg/mL of FL- AF1521^tandem^, or 0.03 μg/mL of FL-eAF1521, and then washed at room temperature with PBST (five washes of 5 minutes each). The detection was done using the ODYSSEY CLx (LI-COR). The sensitivity of FL-AF1521^tandem^ and FL-eAF1521 at given concentrations appeared to be similar for AR-ADPr detection, and FL-AF1521 refers to either probe.

### Antibodies and siRNAs

Commercial antibodies used were anti-Flag M2 (0.75 g/ml, 4°C incubation overnight, Sigma-Aldrich F1804), anti-HA tag (1:1,000, Covance A488-101L, mouse mAb 16b12), anti-Ub (1:20,000, ProteinTech 80992-1-RR), anti-T7 (1:10,000, EMD Millipore 69522), anti-DTX2 (1:3,000, ProteinTech 67209-1-Ig), anti-HUWE1 (0.6 g/ml, ProteinTech 19430-1-AP), anti-TRIP12/ULF (50 ng/ml, abcam AB86220), anti- UBR5/EDD (1mg/ml, Santa Cruz Biotechnology, sc-515494), anti-SPOP (1:10,000, ProteinTech 167501AP), anti-RNF146 (1 g/ml, abcam ab201212), anti-DTX4 (1 g/ml, Aviva Systems Biology ARP68284_P050), anti-Tubulin (1:10,000, Sigma-Aldrich T9028), Fluorescein Antibody DyLight 680 conjugated (1:20,000, Rockland 600-144-096), anti-mono-ADP-ribose (1:1,000, BIO-RAD TZA020), Donkey anti-Human IgG(H+L) Cross Absorbed Secondary Antibody DyLight 800 (1:10,000, Invitrogen SA5-10132), Donkey anti-human IgG(H+L) cross-absorbed secondary antibody DyLight 800 (1:10,000, Invitrogen SA5-10132), Alexa Fluor 680 donkey anti-rabbit IgG(H+L) (1:20,000, Invitrogen A10043), Alexa Fluor 680 donkey anti-mouse IgG(H+L) (1:20,000, Invitrogen A10038), Anti-mouse IgG(H&L) (Goat) antibody DyLight 800 conjugated (1:10,000, Rockland 610-145-002). Lab-made antibodies are anti-AR (1-21 amino acid, 1 g/ml for immunofluorescence), anti-AR(659-667 amino acid, 1 g/ml for Western blot), anti-GST (1mg/ml, T. Parsons’ lab mAb 9D9-F3-F7) and anti-PARP7 (Catalytic domain, 1 g/ml). Flag-AR detection was performed with either anti-AR or M2 antibodies. All of the siRNAs, siCtrl (Am4635), siDTX2 (s223231 & s41519), siDTX1 (s4356), siDTX4 (s23321), siHUWE1 (s19596), siRNF146 (s37822), siSPOP (s15955), siTRIP12 (s17810), siUBR5 (s28025) are from Ambion. Cells were transfected with 20nM siRNA and Lipofectamine RNAiMAX Reagent (Invitrogen 13778-075) for 24hrs, followed by a total of three more days with cell growth and expansion, and drug treatments.

### Drug treatments

Synthetic androgen R1881 (Perkin-Elmer, NLP005005MG; used at 2 nM final concentration in most experiments), natural androgen agonist dihydrotestosterone (DHT, Sigma-Aldrich, D-073), proteasome inhibitor Bortezomib (MedChemExpress, HY-10227; used at 1 M final concentration), PARP7 inhibitor RBN2397 (DC Chemicals, DC31069), PARP14 inhibitor RBN012759 (MedChemExpress, HY-136979), and PARP1/2 inhibitors Veliparib (Selleck Chemical LLC, S100410MG), Olaparib (MedChemExpress, HY-10162), and Talazoparib (MedChemExpress, HY-16106) were used at concentrations and treatment times specified in the figures or figure legends.

### Recombinant protein production

DNA constructs for the GST-DTX2-RD and GST-DTX2-RD^mut^ (S568A, H582A & H594A) with optimized codons were custom synthesized and transformed into E. coli strain *BL21*. Recombinant proteins were induced with 0.4 mM of Isopropyl -D-1-thiogalactopyranoside at 18°C for overnight; cells were harvested and lysed; proteins were purified on a Glutathione-agarose beads (Sigma 4510).

Preparation of recombinant T7-Ub and GST-AF1521^tandem^, and DyLight 800-labeling of the GST- AF1521^tandem^ have been described(*12*).

### In vitro protein binding

The procedures were carried out at 4°C. Cells were extracted in the Buffer A (20 mM Tris-HCl pH 7.5, 100 mM NaCl, 0.5% Triton X-100 (Tx-100), 1 mM PMSF, 2 mM DTT, 5 mM EDTA, 5 µg/ml each of aprotinin/leupeptin/pepstatin (A/L/P) and 1 µM Veliparib with end-over-end rotation for 20 min (0.5 mM ADP-ribose would also be included in the buffer when making immobilized AR that was used for subsequent Ub assays). The extracts were clarified with centrifugation (16,800 x g) for 20 min. The supernatants were subjected to 2.5-4 hrs bindings with anti-Flag M2 magnetic beads (M8823, for immobilizing Flag-AR), or premade Glutathione MagBeads (GenScript L00895-10)/recombinant GST-fusion proteins (3 l packed beads/10 g of protein). The beads were collected under magnetic fields and washed 5-10 times in the Buffer B (20 mM Tris-HCl pH 7.5, 100 mM NaCl, 0.1% Tx-100, 2 mM DTT, 0.1 mM EDTA, 1 µg/ml each of A/L/P and 1 µM Veliparib). The beads were re-suspended in the SDS-loading buffer, heated at 95°C for 5 min followed with SDS-PAGE separation, Western blot analysis and Odyssey CLx (LI-COR) detection.

### Ubiquitylation assays

Ubiquitylation assays were performed at 30°C for 30 min with 1 mM ATP, 100 g/ml Ub (T7-Ub or bovine Ub (Sigma U6253)), 5 g/ml each of UB E1 (R&D Systems E-304) and UB E2 (His-UbcH5C, R&D Systems E2-627), and 20 g/ml GST-DTX2 RD in the buffer E (20 mM Tris-HCl pH 7.5, 50 mM NaCl, 2 mM MgCl2, 1 mM DTT, 1 µg/ml each of A/L/P and 0.1 mg/ml BSA). Preparation of AR as a ubiquitylation substrate was performed by IP as described(*12*), with the addition of an N-Ethylmaleimide treatment step to inhibit trace levels of cell-derived E3 activity. Magnetic M2-Beads, carrying immobilized Flag-AR from PC3-AR(siDTX2) with 6 hrs of 2 nM R1881-treatment and washed thoroughly, were further washed twice with Buffer C (PBS, 1 µg/ml each of A/L/P and 0.1% Tx-100) followed with 5 mM N-Ethylmaleimide/Buffer C treatments at 37°C for 60 min. These beads were washed twice with the assay buffer plus 0.1% Tx-100, once with the assay buffer lacking TX-100, and subsequently used in ubiquitylation assays. Ubiquitylation assays using a chemically synthesized AR peptide (1 M) with ADP-ribosylation on Cys284(*43*) and FITC-labeled on the N-terminus were performed using the same reaction conditions as cell-derived AR. Conjugation products were detected by immunoblotting with anti-Fluorescein (FITC) antibody. NUDT16 reactions were conducted at 30°C for 30 min or at 37°C for 60 min with 2.5, 5, 10, or 25 M NUDT16 in Buffer D (20 mM Tris-HCl pH 7.5, 50 mM NaCl, 5 mM MgCl2, 2 mM DTT, 1 µg/ml each of A/L/P and 0.1 mg/ml BSA). USP2 treatments were carried out at 37°C for 15 min with 1 M His-USP2 CD (catalytic domain, 258-606AA) in the Buffer F (50 mM Tris-HCl pH7.5, 0.5 mM EDTA, 1 mM DTT, 1 g/ml A/L/P and 0.1 mg/ml BSA).

### Immunofluorescence microscopy

Procedures were done at 23°C (room temperature). Cells on coverslips were rinsed with PBS, fixed in 10% Neutral Buffered Formalin for 20 min, rinsed with PBS, permeabilized in PBS/0.2% Tx-100 (PBST) for 20 min, rinsed three times with PBST, blocked in 5% BSA/PBST for 30 min, incubated with 1° Ab/1% BSA/PBST for 2 hrs, washed three times with PBST, incubated with 2° Ab/1% BSA/PBST for 2 hrs, washed three times with PBST, incubated with 2 g/ml 4’,6-diamidino-2-phenylindole/PBS for 15 seconds or more, rinsed with milli-Q water, mounted on slides with ProLong Gold antifade reagent (Invitrogen P36934). Images shown in the figures were acquired by confocal microscopy (Stellaris 8; Leica) equipped with a 100x/1.40 Oil STED White oil immersion objective and Leica software (LAS X).

### Statistical information

All statistical analysis and data visualization was done in R/RStudio.(*69, 70*) Mathematical modeling was performed in MATLAB (version R2023b) using the ODE solver ode23s to solve the ODE system of mass-action equations. Pixel Pearson correlation for images was done in Fiji(*71*) with the JaCoP plugin.(*72*) For the differential expression analysis, p-values were calculated using the two-tailed Wald test. For GSEA, the p-values were calculated using the two-tailed permutation-based testing with 500,000 permutations.

For ORA, the p-values were calculated using the one-tailed hypergeometric test. All adjusted p-values in this study were calculated using the Benjamini-Hochberg (BH) method. For SSE score comparison between models, p-values were calculated using the two-tailed Wilcoxon test. For RT-qPCR relative expression comparisons, p-values were calculated using the two-tailed Welch’s t-test. If the exact p-values aren’t stated, the following symbols were used to indicate significance: ns: p > 0.05; *: p <= 0.05; **: p <= 0.01; ***: p <= 0.001; ****: p <= 0.0001. For qPCR, a bar or a data point is an average of three biological replicates (distinct samples), and error bars represent the standard deviation. For correlation analysis, Pearson correlation was used to calculate a coefficient and p-values (two-tailed). For multivariate Cox regression, p-values were calculated using the Wald test. For the Kaplan-Meier (KM) analysis, p-values were calculated using the two-tailed log-rank test. For RNA-seq and GSEA, we defined p-adj < 0.001 as significant.

## Supplementary Materials

- Supplementary Fig. 1: Gene expression changes in response to androgen and RBN2397 treatment, related to Fig. 1.
- Supplementary Fig. 2: Heatmap showing the module eigengene expression for the 19 modules included in the analysis, related to Fig. 2.
- Supplementary Fig. 3: AR protein levels in cells treated with androgen, RBN2397, and bortezomib, related to Fig. 3.
- Supplementary Fig. 4: AR-Parp7 interactions analyzed by mathematical modeling and confocal microscopy, related to Fig. 4.
- Supplementary Fig. 5: Analysis of AR ADP-ribosylation mutants, related to Fig. 5.
- Supplementary Fig. 6: Mono-ADP-ribose recognition by the DTC domain in DTX2, related to Fig. 6.
- Supplementary Fig. 7: Predictions regarding the effect of DTX2 on Ub conjugation and AR, related to Fig. 7.
- Supplementary Table 1: Model Parameters and Scores
- Supplementary Table 2. AR ADP-ribosylation site mutants used in this study

## Supporting information

Supplementary Fig. 1: Gene expression changes in response to androgen and RBN2397 treatment, related to Fig. 1.

Supplementary Fig. 2: Heatmap showing the module eigengene expression for the 19 modules included in the analysis, related to Fig. 2

Supplementary Fig. 3: AR protein levels in cells treated with androgen, RBN2397, and bortezomib, related to Fig. 3

Supplementary Fig. 4: AR-Parp7 interactions analyzed by mathematical modeling and confocal microscopy, related to Fig. 4

Supplementary Fig. 5: Analysis of AR ADP-ribosylation mutants, related to Fig. 5

Supplementary Fig. 6: Mono-ADP-ribose recognition by the DTC domain in DTX2, related to Fig. 6

Supplementary Fig. 7: Predictions regarding the effect of DTX2 on Ub conjugation and AR, related to Fig. 7

## Acknowledgments

### Funding

NCI award R01CA214872 (B.M.P.)

Saunders Memorial Funding, UVA Comprehensive Cancer Center (B.M.P.) NIH/NCI grant R01CA178393 (A.R.)

NIGMS award R15GM140409 (J.B.K.)

Engineering and Physical Sciences Research Council (EPSRC) studentship through Doctoral Training Partnership EP/W524633/1 (P.A.W.)

Alexander von Humboldt Foundation for a Feodor Lynen Research Fellowship (P.P.)

Author contributions:

Conceptualization, B.M.P., K.W., and C.-S.Y., and A.R.

Methodology, B.M.P., K.W., C.-S.Y., A.R., A.A., J.B.K., P.A.W., and P.P.

Validation, B.M.P., K.W., C.-S.Y., A.A., J.B.K., P.A.W., and P.P.

Formal Analysis, K.W., C.-S.Y., A.R., A.A., J.B.K., P.A.W., P.P., and N.M.D.

Investigation, B.M.P., K.W., C.-S.Y., A.R, P.A.W., P.P., A.A., J.B.K., N.M.D., and J.M.

Resources, D.J.W, S.W. and D.V.F.

Writing – Original Draft, B.M.P, K.W, C.-S.Y., J.B.K., and P.A.W. Writing – Review & Editing, B.M.P., K.W, and C.-S.Y.

Visualization, B.M.P., K.W., C.-S.Y., A.R., A.A., J.B.K., P.A.W., P.P. and N.M.D.

Supervision, B.M.P., A.R., J.B.K., and D.J.W

Funding Acquisition, B.M.P., A.R., J.B.K., D.V.F.

Competing interests: The authors declare no competing interests.

### Data and materials availability

RNA-seq data that support the findings of this study have been deposited in the National Center for Biotechnology Information Gene Expression Omnibus (GEO) and are accessible through the GEO Series accession number GSE272500. All publicly available sequencing data analyzed during this study are available in GEO (ChIP-seq data - GSE84432, RNA-seq data - GSE148397, GSE136272, GSE148397, GSE133876, GSE120660, GSE135879). All other relevant data are available from the corresponding author on request.

