## Supplementary Fig. 1: Gene expression changes in response to androgen and RBN2397 treatment, related to Fig. 1. for "Parp7 generates an ADP-ribosyl degron that controls negative feedback of androgen signaling"

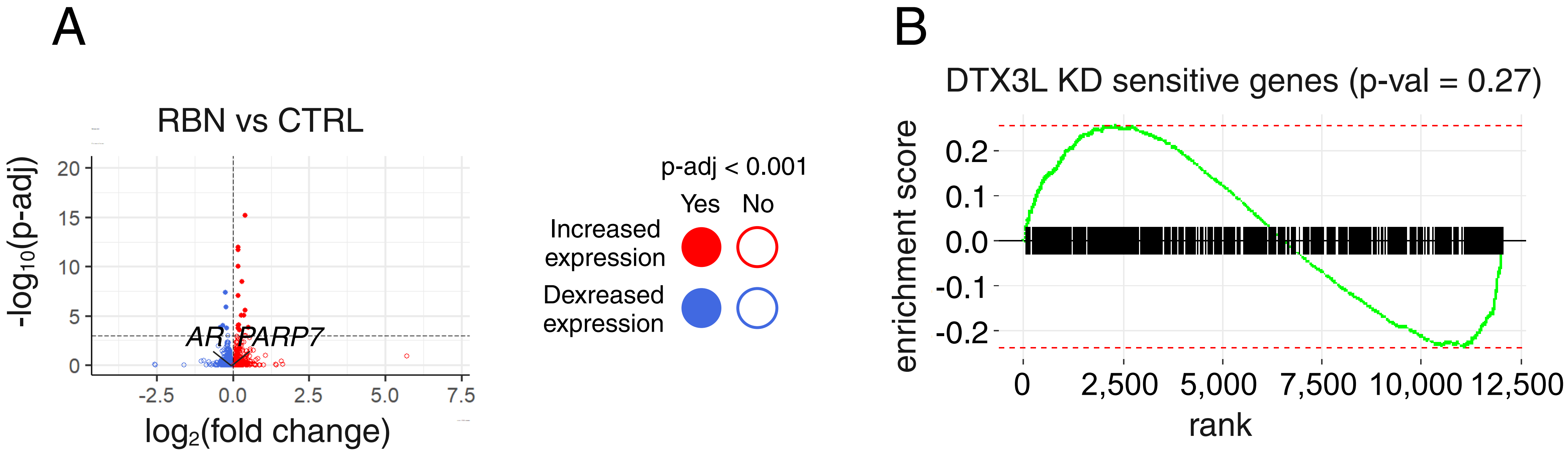


**Supplementary Fig. 1: Gene expression changes in response to androgen and RBN2397 treatment, related to Fig. 1.** A, Volcano plot depicting the differential gene expression analysis in VCaP cell line comparing gene expression between RBN2397-treated and untreated samples (RBN vs CTRL). Each dot represents a gene, with the x-axis showing the $\log_2$ fold change ($Log_2FC$) and the y-axis representing the negative $\log_{10}$ of the adjusted p-value (p-adj). Genes that are significantly upregulated or downregulated are highlighted in red and blue, respectively, based on the preset p-adj (0.001) threshold. B, Enrichment plot depicting the enrichment of genes affected by shDTX3L knockdown in R1881+RBN vs R1881 differentially expressed genes. For the GSEA gene set, we utilized DTX3L KD-sensitive genes from samples treated with R1881, excluding any genes influenced by DTX3L KD under basal conditions. The x-axis in both plots represents a ranked list of R1881+RBN vs. R1881 differentially expressed genes based on log2 fold change, while the y-axis displays the enrichment score values. The green line in the plot shows the trend of enrichment scores, with positive values indicating an increased presence of gene sets toward the top of the ranked list, and negative values indicating an increased presence of gene sets toward the bottom of the ranked list. The analysis rendered insignificant with p-value = 0.27.
