## Supplementary Fig. 2: Heatmap showing the module eigengene expression for the 19 modules included in the analysis, related to Fig. 2 for "Parp7 generates an ADP-ribosyl degron that controls negative feedback of androgen signaling"

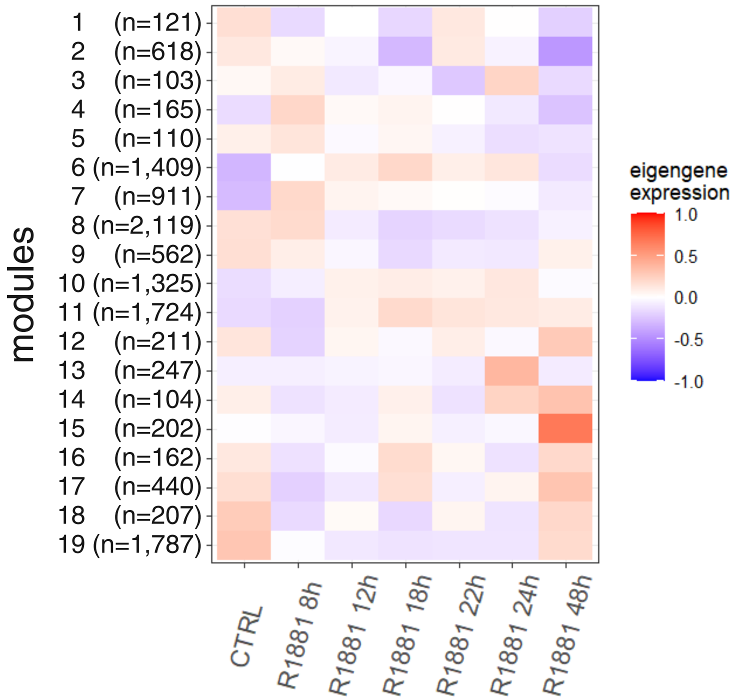


**Supplementary Fig. 2: Heatmap showing the module eigengene expression for the 19 modules included in the analysis, related to Fig. 2.** Module names, together with the number of genes, are presented on the y-axis. The x-axis represents the time points of the R1881 treatment. The representative color gradient used for the heatmap is presented in the figure.
