## Supplementary Fig. 3: AR protein levels in cells treated with androgen, RBN2397, and bortezomib, related to Fig. 3 for "Parp7 generates an ADP-ribosyl degron that controls negative feedback of androgen signaling"

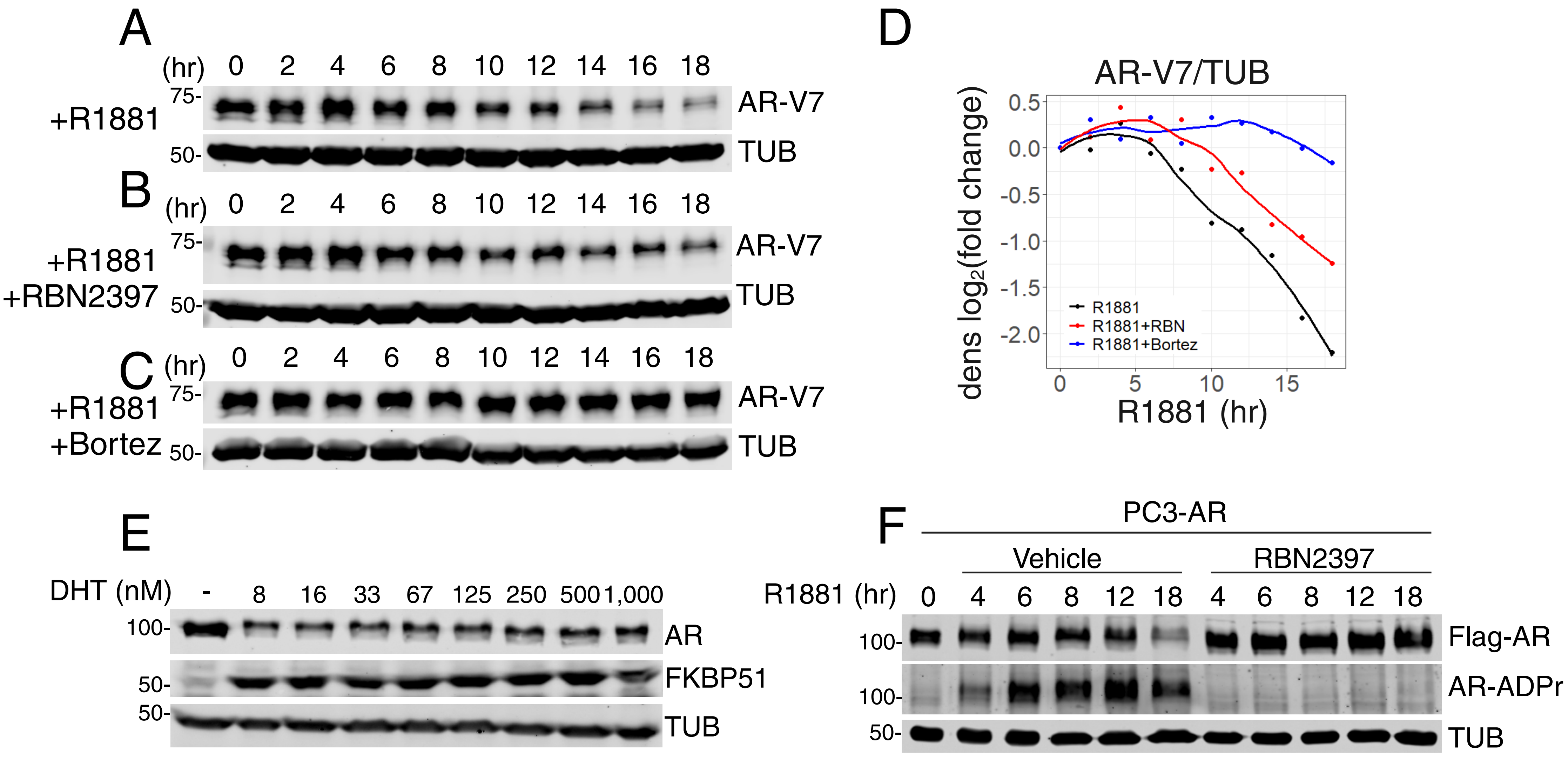


**Supplementary Fig. 3: AR protein levels in cells treated with androgen, RBN2397, and bortezomib, related to Fig. 3.**

A, Immunoblot detection of AR-V7 in VCaP cells subjected to a time course of R1881 treatment. Cells were collected every 2 hours for an 18-hour period (the same experiment as Figure 3A). The Tubulin loading control used for this panel is the same as in Figure 3A as it is part of the same blot.

B, Immunoblot detection of AR-V7 in VCaP cells subjected to a time course of R1881 and RBN2397 cotreatment. Cells were collected every 2 hours for an 18-hour period (the same experiment as Figure 3B).

C, Immunoblot detection of AR-V7 in VCaP cells subjected to a time course of R1881 and Bortezomib (Bortez) cotreatment. Cells were collected every 2 hours for an 18-hour period (the same experiment as Figure 3C).

D, Line plot visualizing the AR-V7 protein density measurements for immunoblots from panels A-C. The y-axis represents the $\log_2$ of the AR-V7/TUB density fold change from 0-hr time point, and the x-axis represents the time of treatment in hours. The curves were fitted using the loess method.

E, Immunoblot detection of AR in VCaP cells treated with a concentration range of DHT for 10 hr.

F, Immunoblot detection of Flag-AR and AR-ADPr (FL-AF1521) in PC3-AR cells treated with R1881 and co-treated with R1881 and RBN2397 for times indicated on the panel.
