## Supplementary Fig. 4: AR-Parp7 interactions analyzed by mathematical modeling and confocal microscopy, related to Fig. 4 for "Parp7 generates an ADP-ribosyl degron that controls negative feedback of androgen signaling"

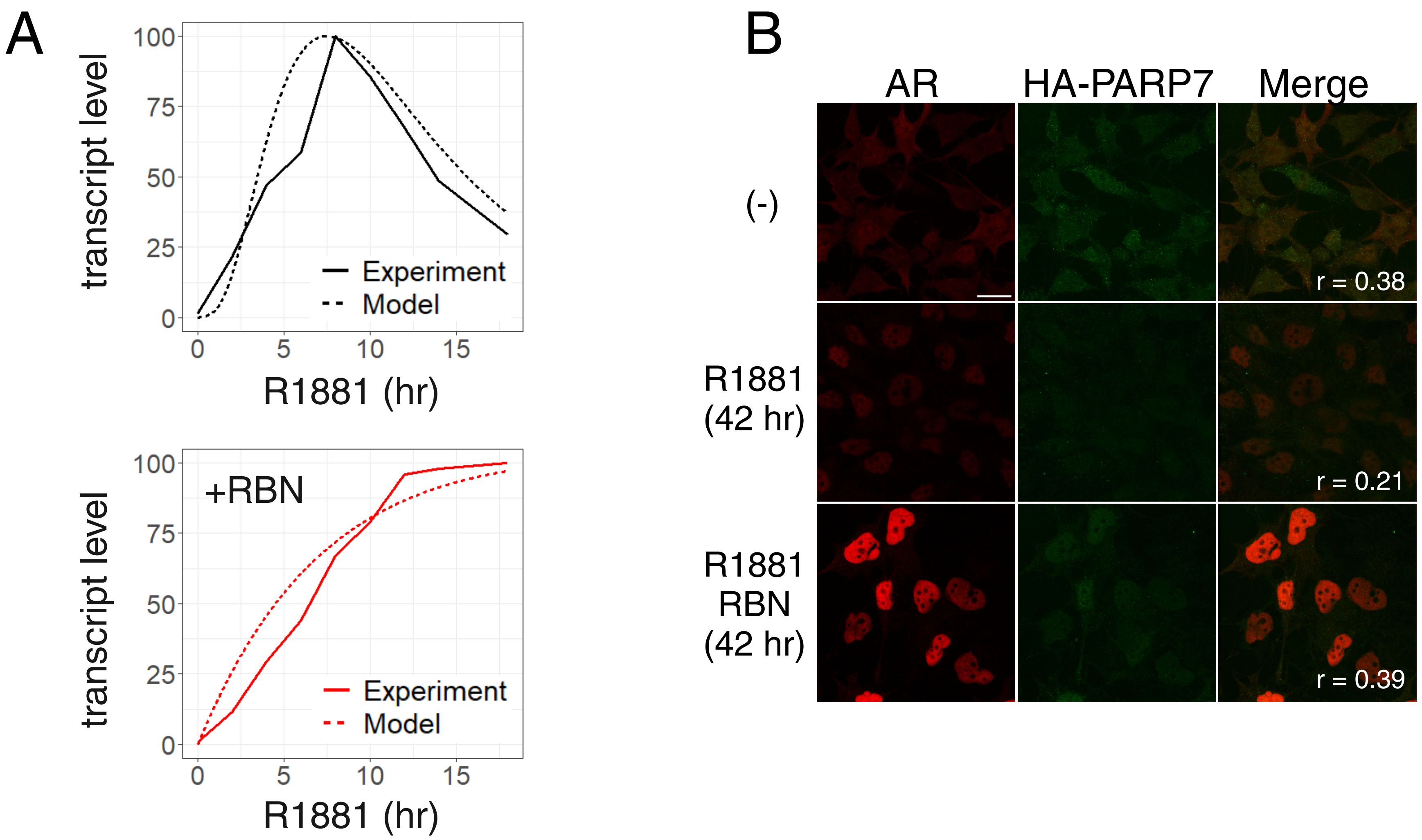


**Supplementary Fig. 4: AR-Parp7 interactions analyzed by mathematical modeling and confocal microscopy, related to Fig. 4.**
A, Plots of experimental transcript for model 2 and simulated transcript during R1881 treatment, with and without RBN2397. Data is normalized to have a maximal value of 100.
B, Confocal microscopy staining of HA-PARP7 and AR in PC3-AR HA-PARP7 cells treated with R1881 and co-treated with R1881 and RBN2397 for times indicated on the panel. The third column shows merged channels, and the Pearson correlation coefficient for pixel colocalization is indicated in the bottom left corner for every condition. A scale bar of 10µm is provided on the bottom right corner of the upper left panel and applies to all panels.
