## Supplementary Fig. 5: Analysis of AR ADP-ribosylation mutants, related to Fig. 5 for "Parp7 generates an ADP-ribosyl degron that controls negative feedback of androgen signaling"

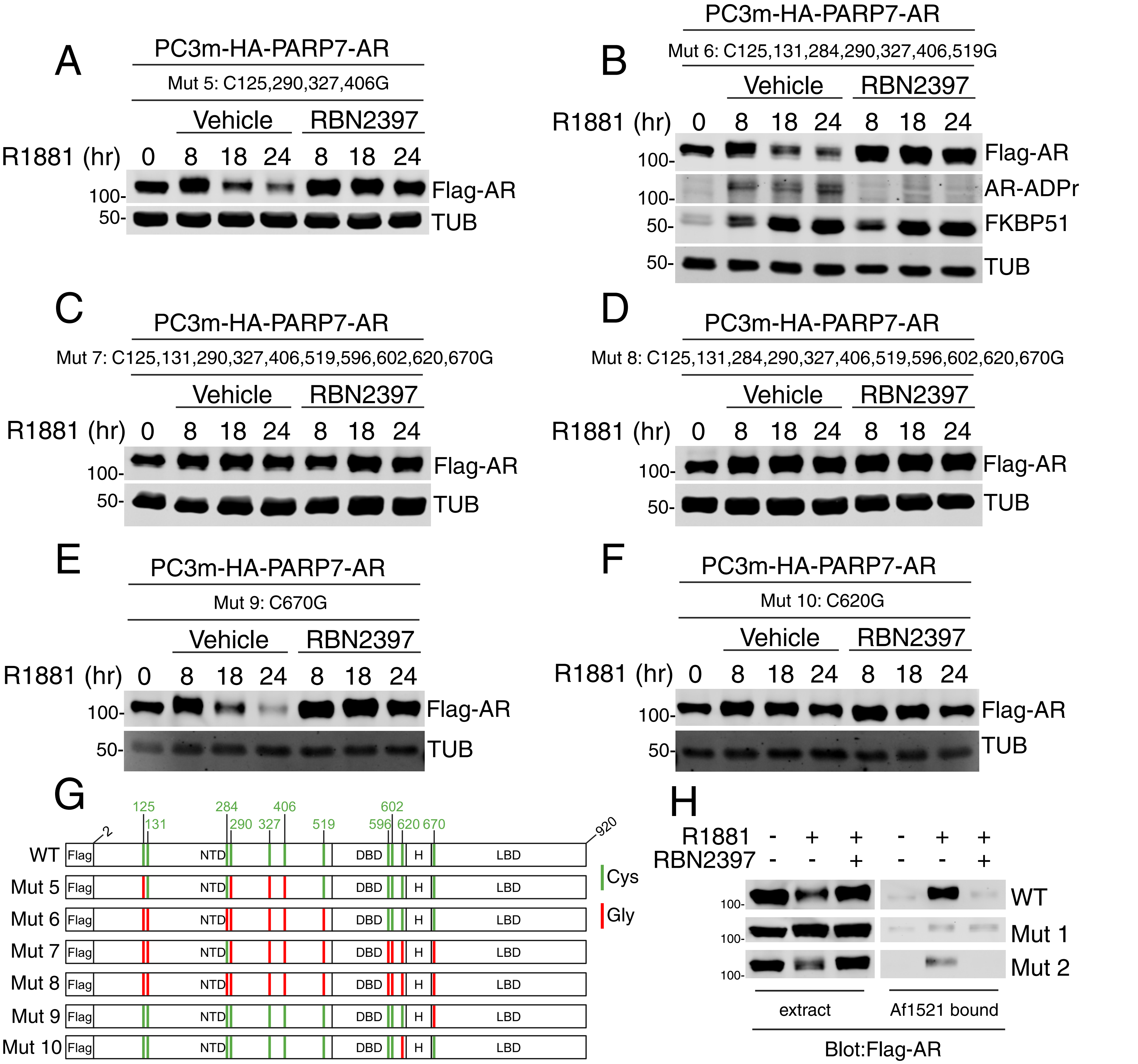


**Supplementary Fig. 5: Analysis of AR ADP-ribosylation mutants, related to Fig. 5.**
A-F, Immunoblot detection of Flag-AR, TUB, AR-ADPr (FL-AF1521), and FKBP51 in PC3m(HA-PARP7/Flag-AR Mutants) cells treated with R1881 or co-treated with R1881 and RBN2397 for times indicated on the panels. A: AR Mut 5 (C125,290,327,406G); B: AR Mut 6 (C125,131,284,290,327,406,519G); C: AR Mut 7 (C125,131,290,327,406,519,596,602,620,670G); D: AR Mut 8 (C125,131,284,290,327,406,519,596,602,620,670G); E: AR Mut 9 (C670G); F: AR Mut 10 (C620G).
G, Diagrams of Flag-AR mutants employed in this figure (Mut 5-10). All of the ADP-ribosyl cysteine sites on AR are marked in green, and the glycine substitutions are marked in red.
H, Immunoblot detection of the Flag-AR protein in PC3m(HA-PARP7/Flag-AR WT, Mut 1 and Mut 2) cell extracts and GST-AF1521tandem bound fractions. Cell extracts from cells treated with different combinations of R1881 and RBN2397 for 18 hr and were combined with magnetic Glutathione/GST-AF1521tandem beads for the enrichment of ADP-ribosylated proteins.
