## Supplementary Fig. 6: Mono-ADP-ribose recognition by the DTC domain in DTX2, related to Fig. 6 for "Parp7 generates an ADP-ribosyl degron that controls negative feedback of androgen signaling"

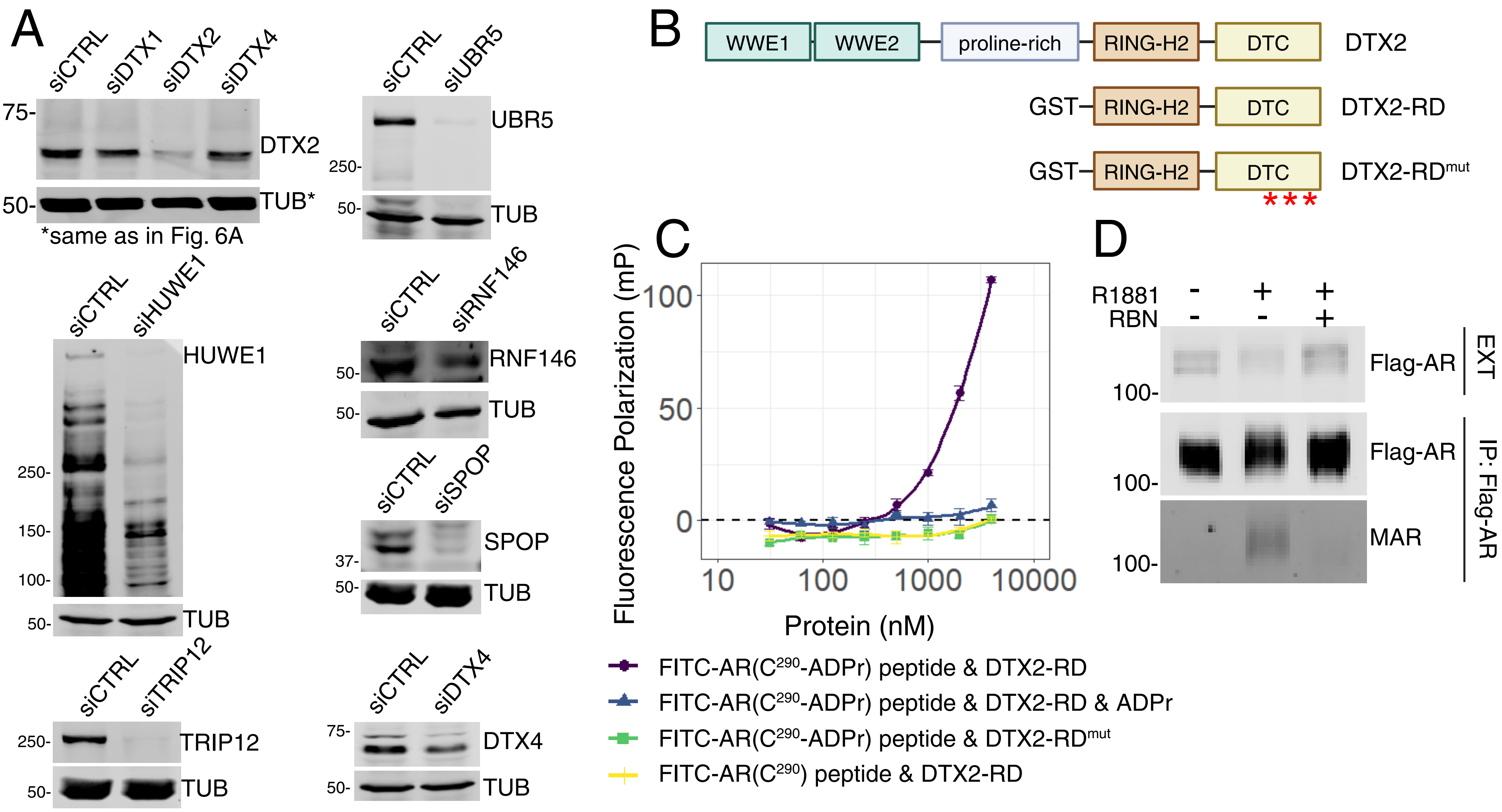


**Supplementary Fig. 6: Mono-ADP-ribose recognition by the DTC domain in DTX2, related to Fig. 6.**
A, Knockdown (siRNA) and immunoblotting of E3 Ub ligases with reader domains, or known roles in nuclear receptor degradation.
B, Diagrams of DTX2 full length, DTX2-RD, and DTX2-RD$^{mut}$. Three loss of function substitutions in the DTC domain of DTX2-RD$^{mut}$ (S568A, H582A, and H594A) are indicated with red asterisk.
C, Line plot presenting the binding measurements of DTX2-RD or DTX2-RD$^{mut}$ to FITC-AR(C290-ADPr) or FITC-AR(C290) peptides by fluorescence polarization. The y-axis represents Fluorescence Polarization (mP) and the x-axis represents the concentration of DTX2-RD or DTX2-RD$^{mut}$. AR peptide sequence: FITC-PLAEC(-/+ADPr)KGSL-OH.
D, Immunoblot detection of the Flag-AR and mono-ADP-ribosylation (MAR) protein in the PC3-AR cell extracts and Flag-AR immunoprecipitation. Cell extracts from cells treated with different combinations of R1881 and RBN2397 for 18 hr and were combined with magnetic anti-Flag M2 beads for the IP.
