## Supplementary Fig. 7: Predictions regarding the effect of DTX2 on Ub conjugation and AR, related to Fig. 7 for "Parp7 generates an ADP-ribosyl degron that controls negative feedback of androgen signaling"

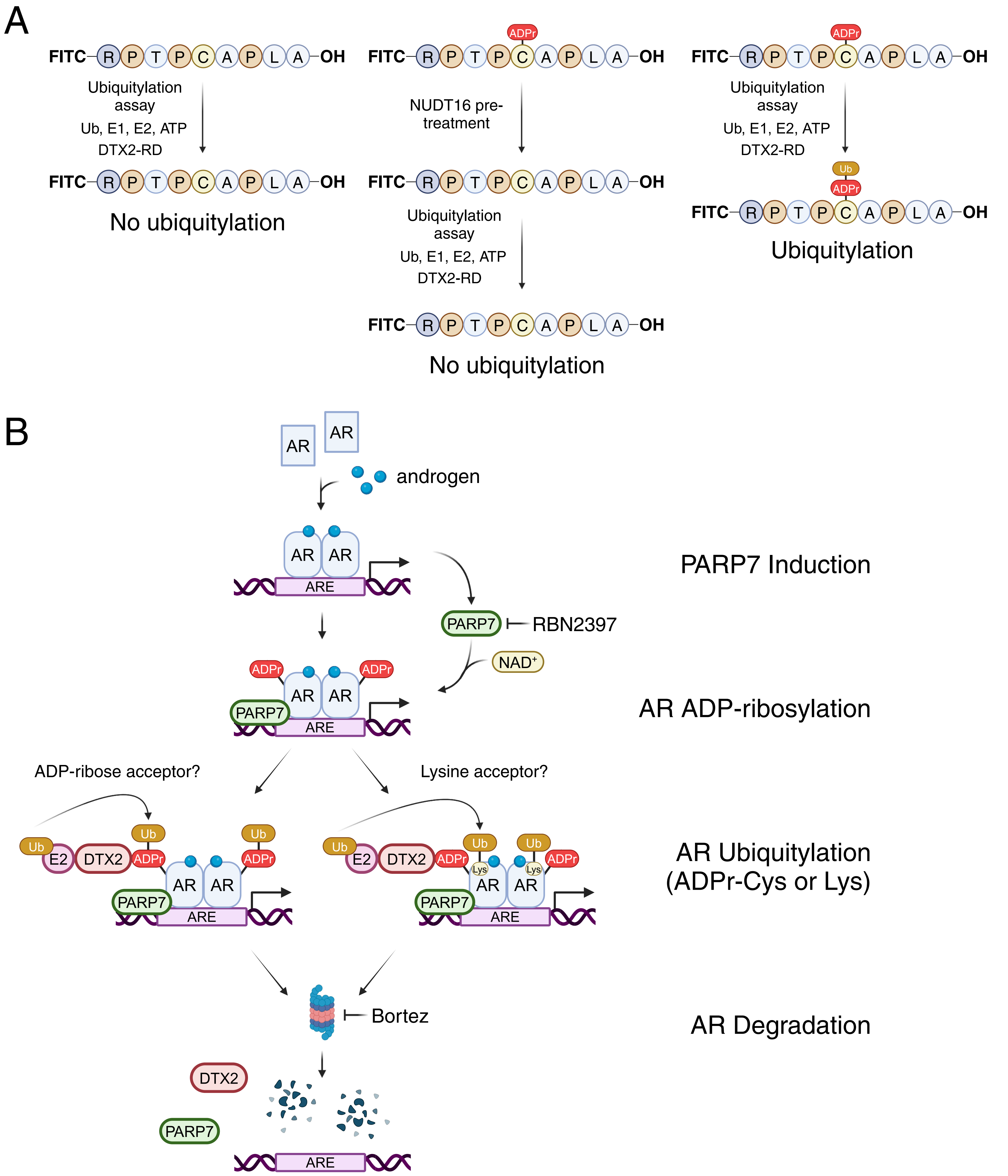


**Supplementary Fig. 7: Predictions regarding the effect of DTX2 on Ub conjugation and AR, related to Fig. 7.**
A, Schematic presenting different molecular outcomes of FITC-AR(C290-ADPr) or FITC-AR(C290) peptides in-vitro ubiquitilation assays from Figure 7A.
B, Schematic presenting the molecular mechanism of negative feedback mediated by DTX2-dependent degradation of ADP-ribosylated AR.
