## Supplementary tables 1&2 for "Parp7 generates an ADP-ribosyl degron that controls negative feedback of androgen signaling"

Supplementary Table 1: Model Parameters

| Parameter | Model 1 | Model 2 | Description of Parameter |
|---|---|---|---|
| $k_1$ | 0.70809 | 0.52759 | ADP-ribosylation of AR driven by PARP7 |
| $k_{1c}$ | 0.16789 | 0.00011 | AR/Promoter complex transcription |
| $k_{1d}$ | 0.50713 | 0.05186 | Degradation of unmodified AR |
| $k_{1r}$ | 0.92896 | 0.83305 | Removal of ADP-Ribosylation from AR |
| $k_{1t}$ | 0.01295 | 0.00794 | Production of AR |
| $k_2$ | 0.85090 | 0.06761 | AR association with the promoter |
| $k_{2c}$ | 0.85968 | 0.56508 | ADP-AR/Promoter transcription |
| $k_{2d}$ | 0.98985 | 0.80955 | Degradation of ADP-AR |
| $k_{2r}$ | 0.78152 | 0.77838 | AR dissociation from the promoter |
| $k_{2t}$ | 0.31458 | 0.79561 | Production of PARP7 influenced by AR |
| $k_3$ | 0.00010 | 0.00013 | ADP-AR association with the promoter |
| $k_{3d1}$ | 0.47086 | 0.85877 | Degradation of PARP7 |
| $k_{3d2}$ | 0.67953 | 0.91768 | Degradation of PARP7 influenced by AR |
| $k_{3r}$ | 0.91374 | 0.74739 | ADP-AR dissociation from the promoter |
| $k_{4d}$ | 0.13421 | 0.13743 | Degradation of Transcript |

| Scores | Model 1 | Model 2 |
|---|---|---|
| BIC | 113.3 | 119 |
| Bayes Weight | 0.95 | 0.05 |

Supplementary Table 2. AR ADP-ribosylation site mutants used in this study

| | **Amino acid substitutions in AR** | **Cys620 status** | **Androgen-induced AR degradation** |
|---|---|---|---|
| **WT** | - | WT | YES |
| **Mut 1** | C125G, C131G, C290G, C327G, C406G, C519G, C620G, C670G | Gly | NO |
| **Mut 2** | C125G, C131G, C290G, C327G, C406G, C519G, C670G | WT | YES |
| **Mut 3** | C125G, C131G, C290G, C327G, C406G, C519G, C620S, C670G | Ser | NO |
| **Mut 4** | C620S | Ser | NO |
| **Mut 5** | C125G, C290G, C327G, C406G | WT | YES |
| **Mut 6** | C125G, C131G, C284G, C290G, C327G, C406G, C519G | WT | YES |
| **Mut 7** | C125G, C131G, C290G, C327G, C406G, C519G, C596G, C602G C620G, C670G | Gly | NO |
| **Mut 8** | C125G, C131G, C284G, C290G, C327G, C406G, C519G, C596G, C602G C620G, C670G | Gly | NO |
| **Mut 9** | C670G | WT | YES |
| **Mut 10** | C620G | Gly | NO |
